# [SNG2], a prion-form of Cut4/ Apc1, enforces non-Mendelian inheritance of heterochromatin silencing defect in Fission Yeast

**DOI:** 10.1101/2022.11.26.517888

**Authors:** Suchita Srivastava, Rudra Narayan Dubey, Poonam Shukla, Jagmohan Singh

## Abstract

Prions represent epigenetic regulator proteins that can self-propagate their structure and confer their misfolded structure and function on normally folded proteins. Like the mammalian prion PrP^Sc^, prions also occur in fungi. While a few prions, like Swi1, affect gene expression, none are shown to affect heterochromatin structure and function. In fission yeast and metazoans, histone methyltransferase Clr4/Suv39 causes H3-Lys9 methylation, which is bound by the chromodomain protein Swi6/HP1 to assemble heterochromatin. Earlier, we showed that *sng2-1* mutation in the Cut4 subunit of Anaphase Promoting Complex abrogates heterochromatin structure due to defective binding and recruitment of Swi6. Here, we demonstrate that the Cut4p forms a non-canonical prion form, designated as [SNG2], which abrogates heterochromatin silencing. [SNG2] exhibits various prion-like properties, e.g., non-Mendelian inheritance, requirement of Hsp proteins for its propagation, *de novo* generation upon *cut4* overexpression, reversible curing by guanidine, cytoplasmic inheritance and formation of infectious protein aggregates, which are converted into monomers upon overexpression of *hsp* genes. Interestingly, [SNG2] prion imparts an enhanced tolerance to stress conditions, supporting its role in promoting cell survival under environmental stress during evolution.

## INTRODUCTION

Following the discovery of prions by Prusiner (1) and their role in transmission of disease traits in humans and cows (2-5), studies in the budding yeast *Saccharomyces cerevisiae* helped to validate the protein-based mechanism of inheritance. Yeast has also provided a genetic system for deciphering the mechanistic details and developing screens for prion treatment. In *S. cerevisiae,* reinvestigation of the phenomena of non-Mendelian inheritance of some mutations reported earlier (6-10), led Wickner to provide evidence in support of protein-based inheritance of the [URE3] ‘prion’ (11). Wickner and others further showed that the [URE3] prion could be generated *de novo* upon overexpression of *URE2* gene in WT but not in *ure2Δ* mutants and was cured by growth in presence of Guanidine hydrochrloride (11-13). Subsequsently, several new prions were discovered, such as, Sup35 with the prion form [PSI^-^](14); [PIN], the prion form of Rnq1, which is required for *de novo* appearance of [PSI^+^] (15, 16) and contains a Q/N-rich domain, that can confer prion function to Sup35 (17); Cyc8, with a Lac^+^ form [OCT^+^] (18) and Swi1 [*SWI^+^*] (19) in *S. cerevisiae* and [Het-s] in Podospora (20). In general, the prion form exhibits loss of function and display protein-based inheritance by virtue its property to convert the normal protein structure into the prion form. As a result, the prion form shows dominance and cytoplasmic inheritance. Because of the dominant nature, the loss of function of the prion form generally leads to dominant negative effect with respect to the WT protein function (21). However, in a few cases the prion form of the protein is mooted to perform normal physiological function or shows a gain of function, such as, Het-s in *Podospora* (20).

So far, only a few cases of prion proteins regulating gene expression have been reported. Among them Swi1, a component of the SWI-SNF yeast chromatin remodeling complex, regulates HO gene expression, Ty insertions *etc*. (19). Similarly, [OCT^+^], the prion form of transcriptional co-repressor Cyc8, causes functional inactivation of the Cyc-Tup1 complex, profoundly affecting global gene expression and cellular phenotypes, including defects in mating, sporulation *etc.* (18). However, there is no report showing the role of prion in regulation of heterochromatin structure and gene silencing. Rather, chromatids assembled into heterochromatin structure have been shown to segregate as Mendelian alleles (22).

Heterochromatin assembly in fission yeast occurs through an evolutionarily conserved pathway (23). According to one model, freshly assembled hyperacetylated histones are first deacetylated at H3-Lys9 position, followed by methylation by the H3-K9-methyltransferase Clr4/Suv39. H3-K9-Me2/Me3 serves as a binding site for the conserved chromodomain protein Swi6/HP1 (24). Further multimerization of Swi6/HP1, together with the cooperative interaction with Clr4, helps to extend the region occupied by the heterochromatin mark (H3-K9-Me2/Me3) and Swi6/HP1 in the flanking regions (25-27).

RNAi machinery also plays a role in silencing, particularly at the stage of establishment of the heterochromatin mark, H3K9-Me, but not in propagation in fission yeast (28). The hallmark of both the heterochromatin and RNAi pathways is their action at the chromatin level: the chromatin structure with its memory acts in a manner similar to a Mendelian allele (22) and does not display cytoplasmic regulation.

We had earlier reported the role of Cut4, the largest subunit of the Anaphase promoting complex (APC; 29), in heterochromatin silencing (30). The mutant allele *sng2-1*/*cut4* shows silencing defect, which exhibited semi-dominance (30), unlike the earlier reported recessive mutants likes *clr1-4*, *swi6*, *etc.* (31-33). Surprisingly, the silencing defect persists even after the *sng2-1* mutation was outcrossed. Further analysis has revealed another property of the encoded protein Cut4: it acts as a prion, which causes loss of silencing at the heterochromatin regions-mating type and centromere. The trait of silencing defect is inherited stably during multiple mitotic divisions and in a non-Mendelian manner during meiosis in a genetic cross with a WT strain. Accordingly, we tested whether this phenotype of silencing defect displays the known characteristics of prions. We show that the silencing defect caused by the putative prion-form of Cut4 is inherited in a non-Mendelian fashion, is cured by treatment of cells with guanidine and overexpression of *hsp104* and *hsp70*, is dependent on *hsp104* for its generation, exhibits dominance and cytoplasmic inheritance and associated with protein aggregation. Importantly, the prion form of Cut4 also shows protein infectivity. Interestingly, the prion derivatives phenocopy the *sng2-1* and *cut4-533* mutants and exhibit enhanced tolerance to various stresses.

## MATERIALS AND METHODS

### Strains and Plasmids

The parental strains used in most of the studies include SPA236 (genotype I: *mat1Msmto REII*Δ*mat2P*::*ura4 ura4D18 leu1-32 ade6-216*) and SPA302 (genotype II: *mat1PΔ17::leu2 REIIΔmat2P::ura4 leu1-32 ura4D18 his2^-^ ade6-210).* The *sng2-1* mutation was transferred to parental strain SPA236 by crossing and checked for silencing effects. Mutations were transferred into the reporter strains by standard genetic crosses. The strains and plasmids used in this study are listed in Table S1 and S2 (Supplementary Data). Media, growth conditions and transformations were as described (34).

### Silencing phenotypes

Normal homothallic strains, denoted as *h^90^* switch at 90% efficiency between Plus and Minus mating type. These cells mate with each other to form zygotes, which undergo sporulation under starvation conditions. The spores have a starchy cell wall which give purple staining with iodine. Thus, cells of efficiently switching *h^90^* strains form colonies that give dark staining with iodine and the phenotype is referred to as spo^+^. In contrast poorly switching mutants, like *swi6^-^*, form colonies that give poor staining with iodine, referred to as spo^-^ phenotype (31, 34).

In contrast, non-switching wt strains with the genotype *Msmto REIIΔ mat2P::ura4*, having a *mat2*-linked *ura4* reporter, have a silent *mat2* locus and give a spo^-^-ura^-^ phenotype. However, in the *swi6^-^* mutant background, they show a spo^+^-ura^+^ phenotype, with colonies giving dark staining with iodine, because the derepression of the silent *mat2Pc* locus, together with Minus information from mat1M locus, triggers meiosis, albeit in haploid cells, producing spores with starchy cell wall.

The loss of silencing can also happen in a diploid state, like in M/M diploid, with genotype: *mat1Msmto REIIΔ mat2P::ura4*/*mat1Msmto REIIΔ mat2P::ura4;* in combination with the homozygous mutation *swi6^-^/swi6^-^,* leading to formation of azygotic asci. This property was exploited in the cytoduction experiment (see Results).

### Mapping of *sng2-1* mutation to *cut4* gene

The *sng2-1* mutation was mapped to *cut4* gene on the basis of the following criteria:

- Complementation of the ts^-^ phenotype of *sng2-1* mutation by *cut4* gene.
- We crossed *sng2-1* and *cut4-533* mutants and dissected 35 tetrads. No ts^+^ segregants were obtained, confirming that *sng2-1* and *cut4-533* mutants were tightly linked.
- We crossed the *sng2-1* mutant with a WT strain and isolated DNA from 5 independent ts^-^ segregants. Sequencing showed that all 5 segregants contained the mutation at 984^th^ position converting codon GTT (encoding Valine) to GCT (encoding Alanine).

### Plasmid Construction

The genes in *S. pombe* were chromosomally deleted and tagged using different heterologous modules. Strains carrying deletion of *swi6*, *clr3*, *hsp104* and *tht1* were constructed using pFA6a-KanMX6 plasmids containing *kan^r^* module. The GFP tagged *cut4* cassette was generated using the tagging vector pFA6a-GFP-KanMX6 (35). The *YFP-FLAG-(His)6* tagged *cut4* (YFH-*cut4*) strain was generated by integrating the YFH- *cut4* at *leu1* site by digesting the plasmid pYFH-*Cut4* with ApaI (procured from Riken DNA Bioresource Center (36).

The *tht1Δ* homozygous diploid was generated by protoplast fusion using inter-alleliccomplementation between *ade6-216* and *ade6-210* alleles (34, 37).

### Determination of the rate of switching of the epigenetic states

We followed the method of Kipling et al (38) to determine the rate of switching. Cells of colonies exhibiting a particular epigenetic state (e.g. Dark or spo^+^, light or spo^-^, ura^+^, ura^w^ or ura^-^) were grown in culture medium and after overnight growth, a known dilution was spread on suitable plate to score the phenotype. Remaining cells were inoculated into fresh medium at cell density of 10^5^ cells/ml and allowed to grow at 30°C for 15-20 generations, after which the same number of cells were spread on the same specific plate as before. After 3-4 days’ growth, number of colonies showing ura^+^ or spo^+^ phenotype were counted. The rate of switching was determined using the formula:

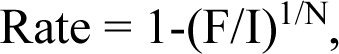

where F is the final percentage of cells having spo^+^ or spo^-^, or ura^+^ or ura^-^ phenotype, I is the initial percentage of cells having spo^+^ or spo^-^, or ura^+^ or ura^-^ phenotype, and N is the number of generations.

### Visualization of prion aggregates and ‘cut’ phenotype

Cells expressing YFP and GFP-tagged *Cut4*p were visualized by confocal microscopy (Leica SP8 Confocal microscope). For visualizing ‘cut’ phenotype, cells were fixed with 70% ethanol, dehydrated, stained with DAPI/PI and visualized under fluorescence microscope. For septum visualization calcofluor staining of the cells was followed by visualization under fluorescence microscope at 433nm (34). For quantitation of cells containing YFP or GFP-Cut4 aggregates, number of cells containing fluorescent foci out of a total of 200 cells were counted.

### Curing by Guanidine hydrochloride

Cells were grown in YEA or selective medium with or without 4mM GuHCl for 5-10 generations. An aliquot of GuHCl-treated cells was washed twice with sterile water and then grown in absence of GuHCl medium for 5 generations. Equal number of cells were taken from each sample, 10-fold serial dilutions were prepared and 2μl of each dilution was spotted on complete, ura^-^ and FOA plates. For quantitative analysis, 50 μl of these cells at OD600 of 0.05 were plated on complete, ura^-^ and FOA plates.

### Curing of prion forms by overexpression of *hsp104* and *hsp70*

*hsp104* of *S. pombe* and *hsp70* gene of *S. cerevisae* were cloned in REP3X vector and transformed into ts^+^-ura^+^ segregants. Equal number of these transformants was plated on leu^-^ ura^-^ plates and ura^+^ cells were counted.

### Thermotolerance assay

Thermotolerance assay was carried out according to Eaglestone e*t al*. (39) with minor changes. Two ura^+^-ts^+^ and two ura^-^-ts^+^ segregants were grown at 30°C to mid log phase in rich medium (OD600 = 0.4) and then transferred to 37°C for 1 h prior to heat treatment. Cells were diluted ∼3.5x 10^3^-fold and transferred to a 52°C shaking water bath. Aliquots were removed at 5 min intervals and stored on ice. From these aliquots 150μl (∼500 cells) were spread in triplicate onto YEA plate and colonies were counted after 4-5 days of growth at 30°C.

### Ethanol tolerance assay

Ethanol tolerance assay was performed according to Eaglestone *et al.* (39). Two ts^+^-ura^+^and two ts^+^-ura^-^ segregants were taken from freshly grown plate, resuspended into 500μl of sterile water and diluted to 1x 10^6^ cells/ml. Stress agar plate was prepared by adding 10% ethanol to YEA medium 24h prior to use. The stress agar plate was allowed to set with the plate tilted, so that there was no stress agar at one end of the plate. Just before use the same volume of YEA devoid of EtOH was poured on to stress agar and allowed to set. Then, 2 μl of cell suspension was spotted on to the stress agar plate along the gradient and sealed with parafilm to prevent evaporation of ethanol and then incubated at 30°C.

### Assay for Heavy metal ion tolerance

Heavy metal tolerance assay was performed according to Yamashita *et al.* (29). Cells of prion and non-prion derivatives were taken from freshly grown plate, resuspended into 500 μl of sterile water and diluted to 1x 10^6^ cells/ml. Chemostress agar plates containing concentrations of CdCl_2_, ranging from 1μM to 20μM, were prepared. 2 μl of cell suspension was spotted on to the stress agar plate and then incubated at 30°C.

### Oxidative stress tolerance

Cells grown to the mid-log phase were subjected to H_2_O_2_ treatment. An overnight culture in 10ml YEL medium was grown up to cell density of *∼*2*×*10^7^ cells/ml (approximately 12^th^ hr of culture growth). Cells were harvested. Test groups containing 0.2mM, 1.0mM or 2.0mM H_2_O_2_ and the control group without H_2_O_2_ were resuspended in 10 ml of YEA medium at the cell density of ∼1×10^7^ cells/ml and incubated ata 30°C in a rotary shaker at 180 rpm for 30 minutes. To calculate cell viability, appropriate dilutions of the cultures were spread on YEA plates. These were incubated at 30°C and number of colonies were counted (40).

### Semi-denaturing Detergent Agarose Gel Electrophoresis

Overnight grown 25ml cultures were harvested, washed with distilled water and centrifuged at 2000rcf for 5min at room temperature. The cells were resuspended in 10 ml of spheroplasting solution having 1.2M D-Sorbitol, 0.5mM MgCl_2_, 20mM Tris.HCl (pH 7.5), 50mM β-mercaptoethanol and 0.5mg/ml Zymolyase and incubated for 30 minutes at 30°C. The cells were checked for spheroplasting under microscope with 1% SDS. The pellet was resuspended in lysis solution consisting of 25mM Tris-HCl (pH 7.5), 15mM MgCl_2_, 15mM EGTA, 1mM DTT, 60mM β- Glycerophosphate, 15mM p-NPP, 0.5mM Sodium Vanadate, 0.1mM Sodium fluoride and 1mM PMSF. Samples were lysed by vortexing at high speed for 5 minutes. The cellular debris was pelleted at 4000 rcf for 5 minutes. The supernatant was removed to a fresh tube. Samples were treated at room temperature for 7 minutes in a 4X sample buffer to give a final concentration of 2% sarkosyl, 5% glycerol, 0.5% TAE, 0.1mg/ml bromophenol blue plus protease inhibitor cocktail (41). Electrophoresis of the samples was performed in 1.5% agarose gel containing 0.1% SDS (42). After electrophoresis, gels were subjected to western blotting with anti-GFP antibody.

### Protein infection

For the protein infection study, a standard protocol was used (43) with some modifications. We prepared the extracts from the prionogenic strains by the spheroplasting method, as reported by Tanaka *et al.* (44) and King and Diaz-Avalos (45). Cell pellet was resuspended in 400μl of lysis buffer, as mentioned above. 50μl glass beads (0.5 mm, Sigma) were added to break cells in a bead beater for five 40 sec cycles with samples being left on ice for 2 min between cycles. Cell debris was removed by centrifugation at 5000 rpm for 5 min at 4°C and the supernatant (‘total protein extract’) was centrifuged at 20,000 x g for 45 min to separate soluble from insoluble fractions. The pellet (insoluble fraction) was then re-suspended in 60 μl of lysis buffer. 20 μl of insoluble fraction was treated with 0.025 units/μl benzonase to digest any nucleic acids present in the sample. Then, 20 ml of exponential phase cell culture of the recipient cells was processed for protein transformation according to Tanaka *et al.* (44) and the transformants were selected on PMA-leu plates. Cell fractionation of normal and derivatives into soluble and pellet fractions was performed according Dulle et al (46).

### Prion domain prediction software

We used the Prion like amino acid composition (PLLAC) software to predict prion forming potential of Cut4 protein (47). This tool predicts Prion-like domains (PrLDs) based on experimentally characterized prion proteins, by identifying Q/N-rich regions in the protein sequence. The protein sequence is modelled as Markov process (HMM), and the state can either be PrLD or background. As part of the analysis the server also identifies regions with IDR using Foldindex and PAPA software.

## RESULTS

### *sng2-1*, a *cut4* allele with dominant negative effect on silencing, exhibits two epigenetic states

We reported earlier that a ts mutant, named *sng2-1*, that grew well at 25°C and 30°C but failed to grow at 36°C, exhibited silencing defect at the mating type and centromere loci in fission yeast (30). The mutation was mapped to *cut4* gene (Materials and Methods) In the strain background *leu1^-^Msmto REIIΔmat2::ura4 ura4D18* (strain name SPA236; genotype I) with *Msmto* representing the unswitchable Minus allele at the *mat1* locus, *REIIΔ* representing the deletion of the repression element *REII*, which is inked to the *mat2P* locus and *ura4* representing the *ura4* reporter gene inserted at centromere-distal location with respect to the *mat2P* locus) (32, 33) (**Figure 1A**), the *sng2-1* mutant exhibited two metastable states with distinct phenotypes: i) D (Dark or spo^+^, which give dark purple staining with iodine, due to loss of silencing and termed as DSPR) and ura^+^ (representing strong growth on plates lacking uracil) due to derepression of the *ura4* reporter and ii) L (light, spo^-^, which, like the parent strain SPA236, give no staining with iodine, and termed as LSPR) and ura^w^ (representing weaker growth on uracil lacking plates) due to partially derepressed *ura4* reporter (34; **Figure 1C**). In contrast, in the strain background with the genotype: *leu1^-^mat1PΔ17::LEU2 leu1-32 REIIΔ mat2::ura4*, *his2^-^* (denoted as genotype II), which is similar to the genotype I except the *mat1* locus harbors the Plus allele along with a linked deletion called Δ17 and insertion of the *LEU2* reporter, which are located centromere distally and linked to *his2^-^* marker (48), the *sng2-1* mutant exhibits spo^-^-ura^+^ phenotype, **Figure 1B**). A reciprocal growth level was observed on FOA plates: ura^+^ cells grow poorly on FOA plates (FOA^s^), while ura^-^ cells grow well on FOA plates (FOA^r^) (**Figure 1C**). This was accompanied in the strain having the genotype I by strong expression of the silent copy transcript *mat2Pc* in the DSPR and weaker expression in the LSPR, but no expression in the WT parent (**Figure 1D**). The two states DSPR and LSPR were metastable, switching to the opposite state at the rate of ∼10^-4^/generation during mitosis (**Supplementary Figure S1B**). Interestingly, the spo^+^ phenotype of the *sng2-1* mutant in the genotype I could not be complemented by the *cut4* gene on high copy vector and only partially by *cut4* gene on integrating vector, as indicated by iodine staining of the transformant colonies (**Figure 1E**). In contrast, the parent WT strain with genotype I did not give any iodine staining (referred to as spo^-^ phenotype; not shown). This indicates that the silencing defect of the *sng2- 1* mutant was dominant or semi-dominant, unlike the recessive defect in the canonical silencing mutants *swi6*, *clr1-4*, *etc*. (31-33; see below).

**Figure 1.**
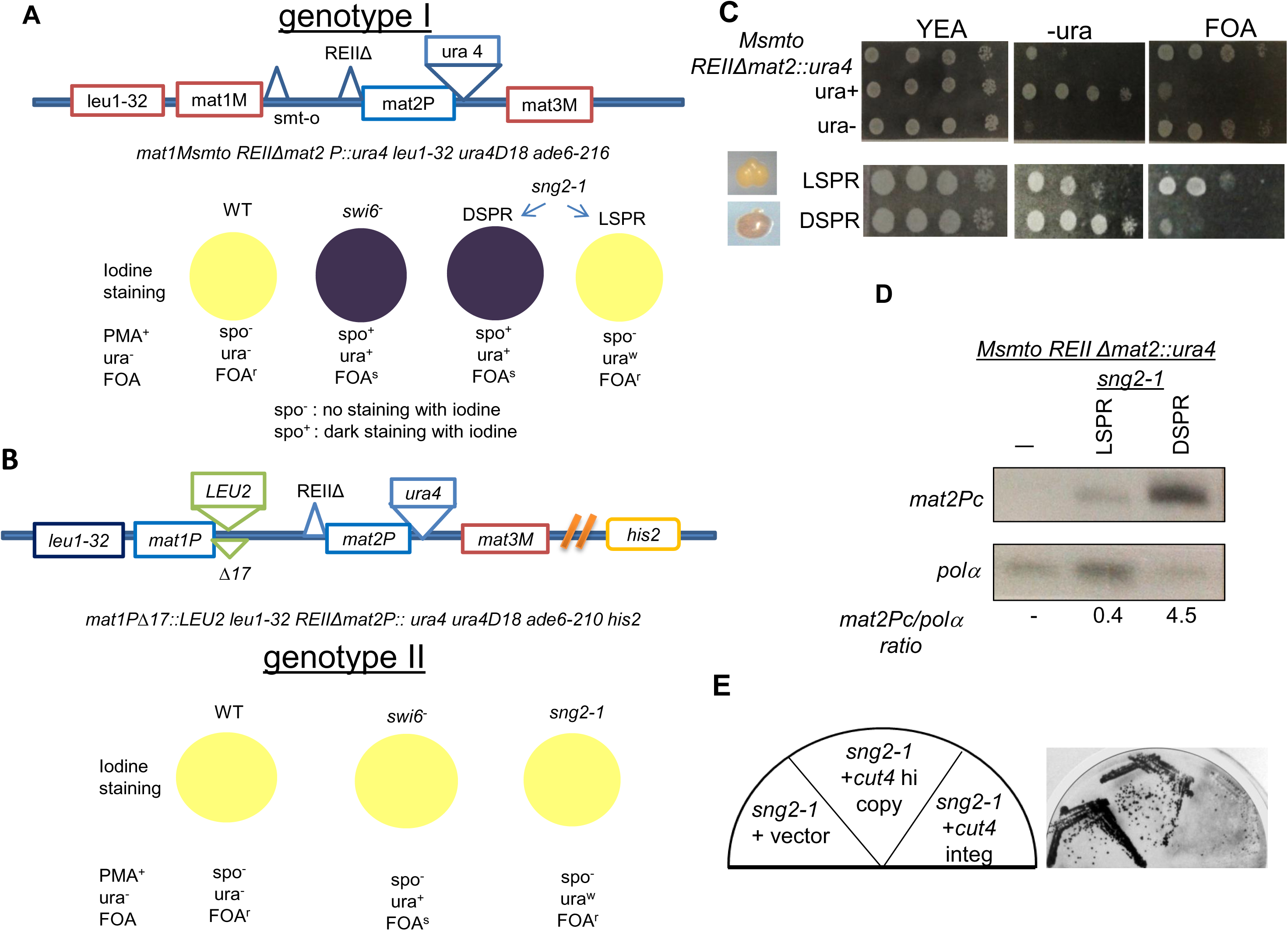
Two alternative silencing defective states displayed by the *sng2-1*/*cut4* mutant.

**(A)** shows the genotype I, denoted as *leu1-32 mat1Msmto REIIΔmat2P::ura4* along with the associated phenotypes in the wt, *swi6^-^* and *sng2-1* background, giving the respective iodine staining phenotypes of the colonies on minimal media (PMA^+^): spo^-^, as light staining and spo^+^ representing dark staining. Also shown are the respective phenotypes of growth on media lacking uracil as ura^-^ or ura^+^, FOA^r^ or FOA^s^. **(B)** Genotype II, denoted as *leu1-32 mat1PΔ17::LEU2, REIIΔmat2P::ura4*, with a linked *his2^-^* marker. Also shown are the phenotypes associated with wt, *swi6^-^* and *sng2-1* backgrounds. **(C)** Dilution spotting assay to assess the level of growth of the control strain with genotype I along with control ura^+^ and ura^-^ strains, on indicated plates. Also shown are the spo^-^ (LSPR) and spo^+^ (DSPR) derivatives of the *sng2-1* mutant with genotype I. **(D)** RT PCR analysis to monitor the level of the *polα* and *mat2Pc* transcripts in wt (-) and LSPR and DSPR derivatives of the *sng2-1* mutant. **(E)** Iodine phenotypes of the *sng2-1* mutant in background of genotype I transformed with vector, *cut4* gene in high copy vector or an integrating vector.

### The spo^+^- ura^+^ phenotype persists after outcrossing the *sng2-1* mutation

To further understand the dominant nature of the silencing defect, the *sng2-1* mutant in the genotype I with spo^+^ and ura^+^ phenotype (DSPR) was crossed with a WT strain having genotype II (**Figure 2**). WT strains in both genotypes I and II give spo^-^-ura^-^ phenotype, while the *sng2-1* mutant gives two inter-switching phenotypes of spo^+^-ura^+^ (DSPR) and spo^-^-ura^w^ (LSPR) in genotype I, as mentioned above, and spo^-^-ura^+^ phenotype in genotype II background (**Figure 1A, 1B**) (+ and– represented the strong and negative phenotypes, respectively; see Materials and Methods).

**Figure 2.**
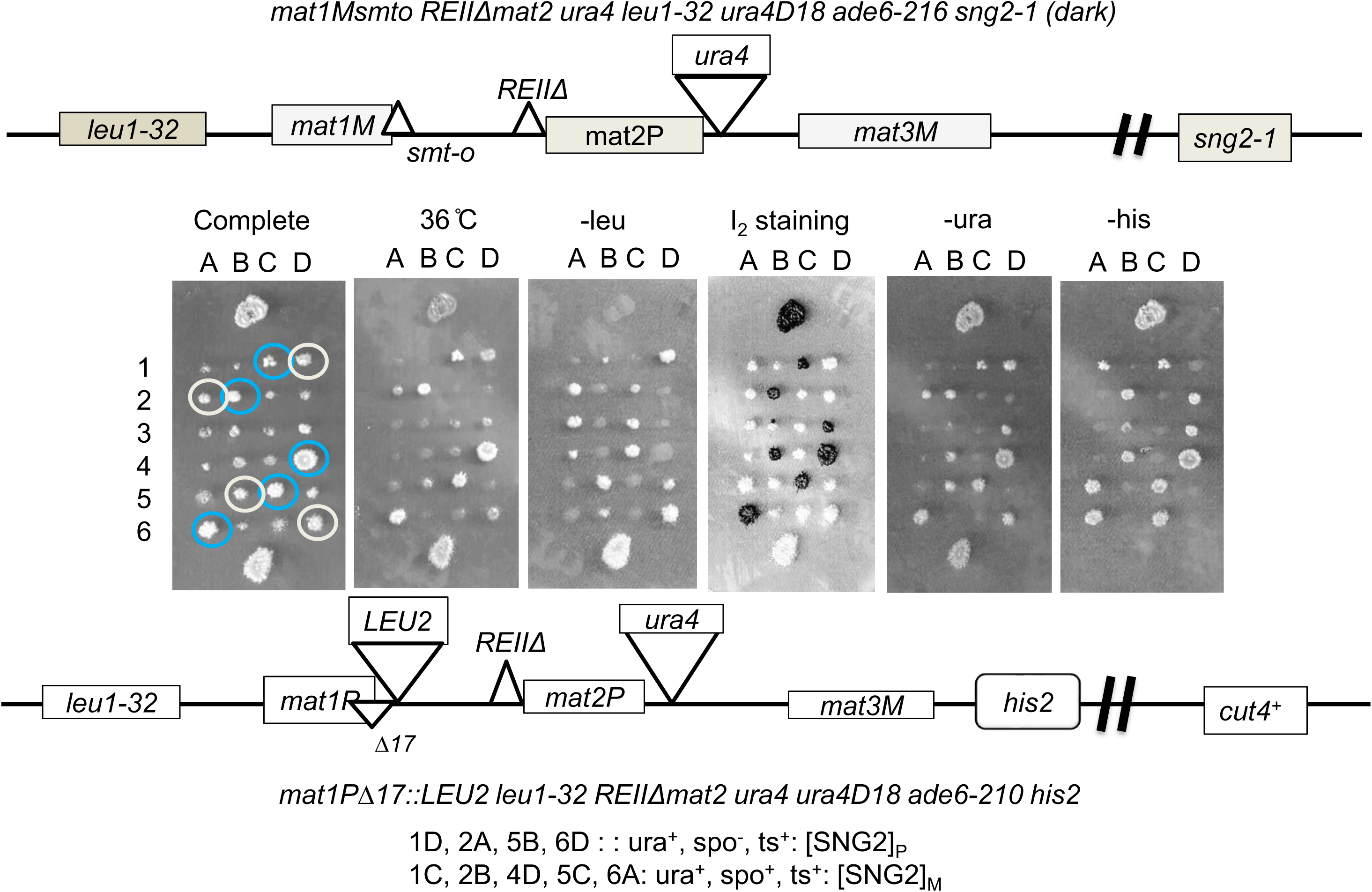
Persistence of the derepressed phenotype of the *sng2-1* mutation after outcrossing. Tetrad analysis shows the cross between the DSPR strain with genotype I (*leu1-32 mat1Msmto REIIΔmat2P::ura4, ura4D18, sng2-1*) with a WT strain having genotype II (*leu1-32 mat1PΔ17::LEU2, leu1-32, REIIΔmat2P::ura4, ura4D18, his2^-^*). Segregants listed as putative “[SNG2]_M_” prion form are circled in blue and those exhibiting the putative prion form “[SNG2]_P_^”^ are circles in white.

Surprisingly, tetrad analysis of the cross between the strain DSPR and the wt strain having genotype II (with ts^+^, spo^-^, ura^-^ phenotype) showed that several ts^+^ segregants with genotype I (segregants 1C, 2B, 4D, 5C and 6A, in which the *sng2- 1* mutation was outcrossed) showed spo^+^-ura^+^ phenotype (**Figure 2**). Interestingly, several ts^+^ segregants (1D, 2A, 5B, 6D) with genotype II also showed ura^+^ phenotype (**Figure 2**). These results indicate that the phenotype engendered by the *sng2-1* mutation persisted even after it was outcrossed. We denoted the leu^-^-ts^+^- spo^+^-ura^+^ segregants with genotype I as putative “[SNG2]_M_” prion-forms and those with genotype II with phenotype of leu^+^-spo^-^-ura^+^-his^-^ as putative “[SNG2]_P_^”^ prion-forms, where M and P represent the mating types *mat1M* and *mat1P*, respectively, the latter being linked to the *LEU2* and *his2*^-^ markers (**Figure 2, 3A**).

**Figure 3.**
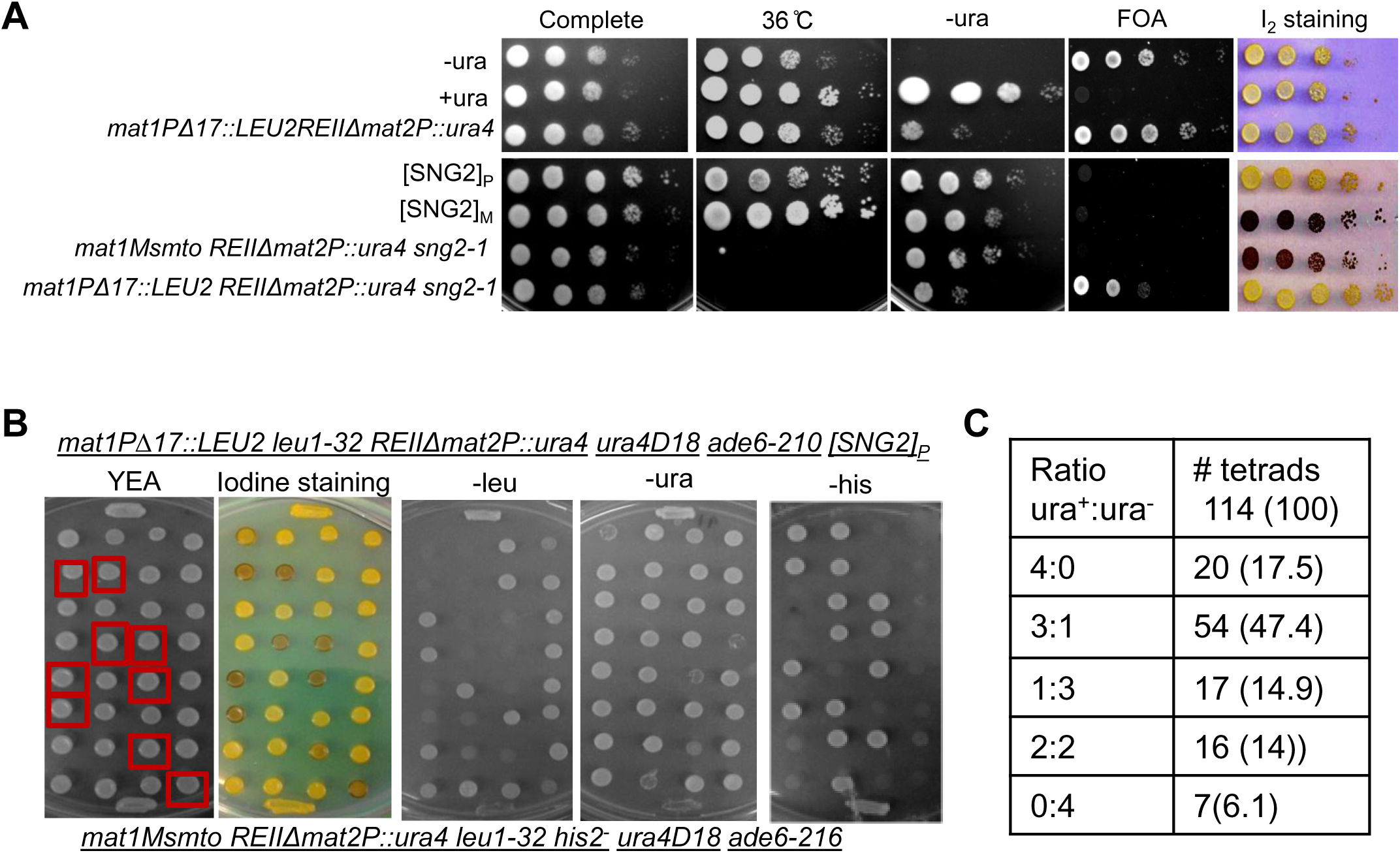
Non-Mendelian segregation of the putative [SNG2] prion-form. **(A)** Phenotypes of the wt, *sng2-1* mutant and the putative prion derivatives, [SNG2]_M_ and [SNG2]_P_, in the genetic backgrounds I and II, respectively, was monitored by spotting assay on indicated plates. **(B)** Tetrads derived from a cross of [SNG]_P_ strain showing the spo^-^-ura^+^ phenotype (top), and a WT strain (genotype I) showing the spo^-^-ura^-^ phenotype (bottom), were spotted on YEA and then replica plated to different plates, as indicated. Red boxed segregants represent the putative “[SNG2]_M_” prion-form. **(C)** Segregation pattern of ura^+^ phenotypes of the *mat2P::ura4* locus in tetrad analysis as shown in **(B)**.

We further tested whether the states “[SNG2]_P_” and “[SNG2]_M_” exhibit the known characteristics of prions.

### Putative prion form “[SNG2]^”^ is inherited in a non-Mendelian manner

To further investigate the possibility that the ura^+^ segregants display the properties of prions, we tested the pattern of inheritance of the ura^+^ phenotype of the“[SNG2]_P_” prion form (from **Figure 2**; Genotype: *leu1-32 mat1PΔ17::LEU2, REIIΔmat2P::ura4*) by back-crossing it with WT cells of strain SPA236 (Genotype: *leu1-32 mat1Msmto, REIIΔmat2P::ura4*). Interestingly, while both the parent strains show spo^-^ phenotype, we observed several spo^+^ segregants (indicated by boxes, **Figure 3B**). In addition, we observed the ura^+^: ura^-^ segregation pattern deviating from the Mendelian ratio of 2:2, with a majority of tetrads (∼86%) showing a non-Mendelian segregation pattern (4:0, 3:1, 1:3 and 0:4) with only 14% tetrads showing the 2:2 Mendelian segregation pattern (See -ura panel **Figure 3B, Figure 3C**). This pattern of inheritance is similar to that displayed by the [URE3] prion (11). These results confirm the non-Mendelian pattern of segregation of the derepressed state, which was engendered by the *sng2-1*/*cut4^-^* mutation and persisted even after the mutation was outcrossed.

More tellingly, we found that among the 104 ts^+^ segregants of a back cross of *sng2-1* mutant with a wt strain, 64 (61.5%) were ura^+^ (representing the derepressed *mat2P-* linked *ura4* locus), as against 0% expected by Mendelian segregation **(Figure S1A)**. We infer that WT segregants with *mat1* Plus background (genotype II) also gave ura^+^ phenotype, unlike the WT parent, which gives spo^-^, ura^-^ phenotype **(Figure S1A)**. Lastly, the spo^+^ phenotype was observed among 47% and 34% segregants among the ts^-^ and ts^+^ segregants, respectively, while the expected level according to the Mendelian segregation would be 56% and 0%, respectively. Overall, these results indicate a non-Mendelian segregation of the *sng2-1*- engendered silencing phenotypes. Furthermore, these states of derepression (spo^+^ and/or or ura^+^) or repression (spo^-^ and/or or ura^-^) exhibited a high level of stability as they switched to the opposite state at a low rate, ranging from 10^-3^ to 10^-6^/generation **(Figure S1B)**. Thus, these states represent metastable states of silencing, as reported by Grewal and Klar (22) although their pattern of inheritance is non-Mendelian (see below).

In contrast, a cross between wt strains with genotype I and II did not yield any tetrads with the spo^+^- ura^+^ phenotype **(Figure S2A)**. Furthermore, in a cross of *clr3Δ*:: kan^r^ mutant in genotype I having spo^+^-ura^+^ phenotype with wt genotype II strain, only *clr3Δ*(G418^r^) segregants showed spo^+^-ura^+^ phenotype, while *clr3^+^* (G418^s^) segregants did not **(Figure S2B)**. These results show that the phenotype of canonical heterochromatin mutants is inherited in a Mendelian pattern. This is diametrically different from the propagation and inheritance of the silencing defect elicited by the *sng2-1*/*cut4^-^* mutation.

Prions have been shown to form protein aggregates (54). To investigate the occurrence of Cut4 as amyloid we repeated the cross shown in **Figure 3B** except that the one of the parent strains (parent I) was constructed by tagging strain having genotype I (SPA236) with YFH- triple tagged *cut4^+^* gene at the *leu1^-^* locus, which is linked to *mat1*. This strain showed a faint diffused YFP fluorescence (**Figure 4B, parent 1 in 4A)**. As shown earlier in **Figure 3B**, segregants with varying phenotypes were observed: spo^+^-ura^+^, spo^-^-ura^+^, spo^+^-ura^w^, (where w represents weak phenotype) in the tetrads and non-Mendelian segregation pattern of ura^+^ phenotype was observed (4ura^+^:0 ura^-^, 3ura^+^: 1 ura^-^) (**Figure 4A**). Consistent with the expectation of Cut4 existing as a prion, while the parent 1 gave faint fluorescence and the second parent (lacking the YFP- cu4 gene) none, the spo^+^-ura^+^ segregants 2B and 3B showed punctuate pattern of YFP fluorescence apparently localized to the cytoplasm (**Figure 4B**). On the other hand, the segregant 1A with spo^-^- ura^-^ phenotype showed faint fluorescence of YFP-Cut4, similar to the parent strain with genotype II (parent 1, **Figure 4B**).

**Figure 4.**
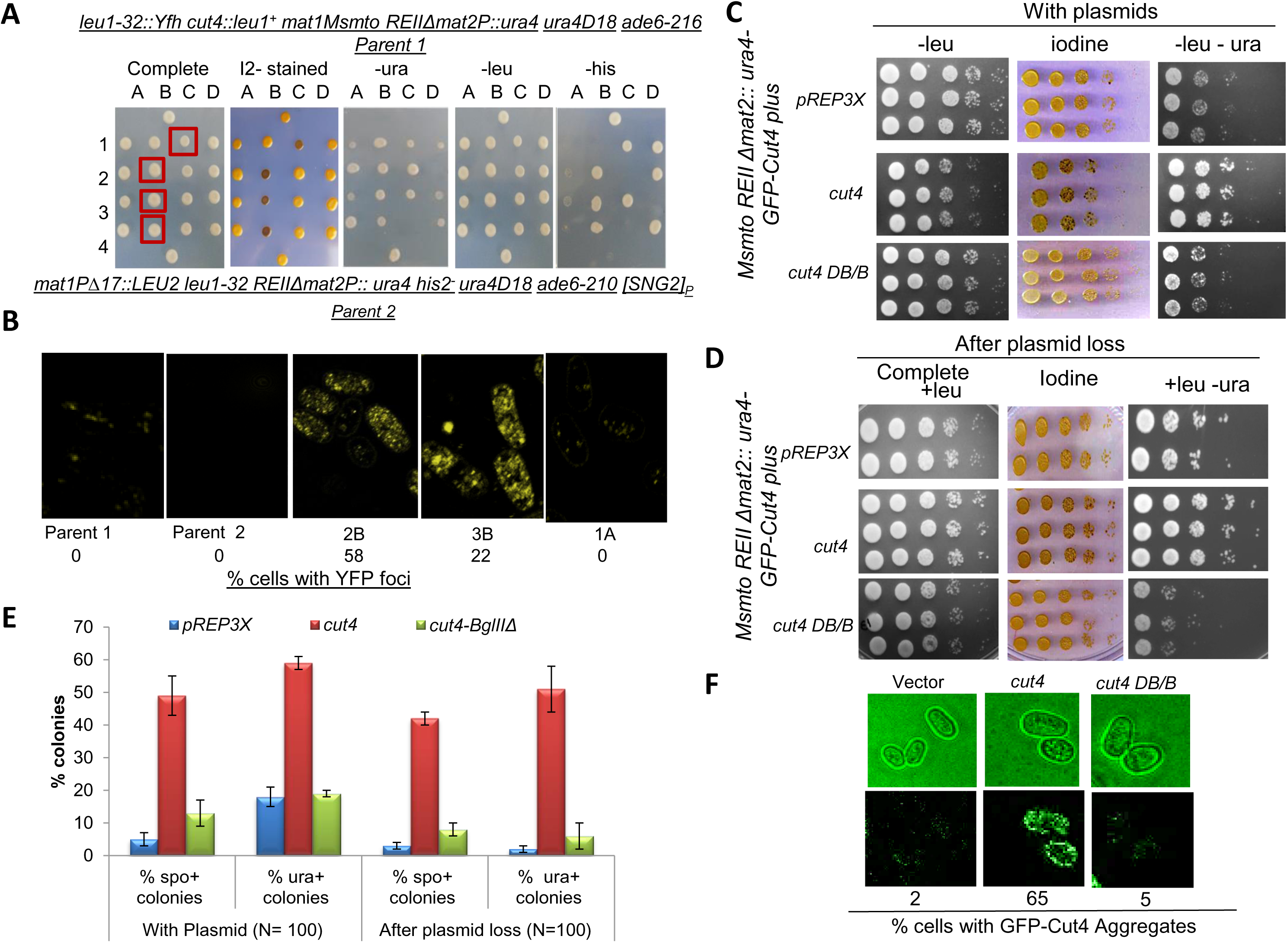
Aggregate formation by Fluorescent-tagged Cut4 upon prion conversion. **(A)** Tetrads derived from a cross between a WT strain of Minus mating type (genotype I) containing an ectopically integrated YFP-Cut4 (top) and an [SNG2]_P_ strain with genotype II (bottom). Orange boxed segregants having the putative “[SNG2]_M_” phenotype are shown. **(B)** Confocal microscopic analysis of selected segregants from tetrads analysis shown in (A) showing YFP-Cut4 aggregation. **(C-F)** *de novo* generation of prion form [SNG2]_M_ upon over expression of *cut4.* **(C)** Serial dilution spotting assay to show growth on plates lacking uracil and iodine staining phenotype, for transformants of WT strain (*leu1-32 Msmto REIIΔmat2ura4*) containing high copy empty vector (pREP3X), high copy vector containing full length *cut4* gene or *cut4* gene having an internal deletion *BglIIΔ* (*cut4*DB/B). **(D)** Serial dilution spotting assay showing the persistence of elevated level of ura^+^ phenotype in WT strain after plasmid loss of the transformants, as shown in **(C)**. **(E)** Quantitation of spo^+^ and ura^+^ colonies upon ectopic expression of vector, *cut4* and *cut*4DB/B on high copy vector and upon loss of vectors, as shown in A and B, respectively. Data was represented as SD. **(F)** Confocal microscopy pictures of cells of WT strain harboring integrated GFP tagged copy of *cut4* gene upon transformation with different vectors. Numbers denote the percentage of cells containing GFP-Cut4 aggregates.

### *de novo* generation of [SNG2] upon overexpression of *cut4* and its non-Mendelian inheritance pattern

It has been shown that high levels of a prionogenic protein can enhance the rate of its conversion to prion form (11). Accordingly, we transformed the strain SPA236 having genotype I (*leu1^-^ mat1Msmto REIIΔmat2P::ura4*) in which the endogenous copy of the *cut4* gene is tagged with GFP, with empty vector, high copy vector containing *cut4* gene or truncated *cut4* gene carrying deletion of its internal *BglII-BglII* spanning region (*DB/B*), which essentially produces a Cut4 protein truncated after 10 amino acids beyond the *BglII* site. We compared several independent transformants for the iodine staining and *ura4* expression phenotypes. We found that the majority of transformants overexpressing *cut4* gene showed elevated level of spo^+^ (∼50%) and ura^+^ (∼60%) phenotypes (**Figure 4C**). We further tested whether, as in the case of prions reported earlier, the spo^+^ and ura^+^ phenotype of the transformants persisted after plasmid loss. We subjected 3 independent colonies (spo^+^ colonies in case of *cut4* overexpression shown in **Figure 4C, 4E**) to plasmid loss and repeated the spotting assay. Results showed that the derivatives of spo^+^- ura^+^ continued to exhibit spo^+^-ura^+^ phenotype after loss of the *cut4* plasmid (**Figure 4D, 4E**). In contrast, overexpression of *cut4* on a low copy vector had no effect (**Supplementary Figure 3A**). Generation of prion form upon overexpression of the gene has been shown to be dependent on the presence of a WT copy of the gene (11). However, this experiment could not be performed as *cut4* gene is essential for viability.

To test the mode of inheritance of the spo^+^-ura^+^ phenotype, we first subjected the spo^+^-ura^+^ transformants of *cut4* on high copy vector to plasmid loss. One such spo^+^- ura^+^ derivative obtained after plasmid loss, tentatively designated as [SNG2]°, was crossed with the strain SPA302 with genotype II (*mat1PΔ17::LEU2 REIIΔ mat2::ura4*). Tetrad analysis showed that the ura^+^ phenotype segregated in a non-Mendelian pattern with 74% tetrads showing the 4:0, 3:1 and 1:3 segregation pattern (Supplementary Figure S3B)

To directly visualize the Cut4 protein aggregates, we carried out confocal microscopy of the spo^+^-ura^+^ transformants of cells containing GFP-tagged *cut4* gene. Results showed that the transformants obtained with the *cut4* on high copy vector contained aggregates of GFP-Cut4, while those carrying the control vector or truncated *cut4* gene (*DB/B*) did not (**Figure 4F**). As shown earlier (**Figure 3E**) the aggregates seem to be localized throughout the cells (**Figure 4F**). The effect was dependent on expression of full-length *cut4* gene as the vector containing truncated copy of *cut4* (*DB/B*) had no effect (**Figure 4F).**

### Curing of [SNG2] by guanidine and [SNG2] propagation by *hsp104*

As protein folding-refolding dynamics underlies the function of proteins, generation of prion form is generally found to be dependent on components of the protein refolding pathway (49, 50), although some conflicting results show lack of such dependence (51).

It has been shown that growth in low concentration of guanidine (5mM) can cure the prion phenotype; Hsp104 is inactivated by low concentration of guanidine, which leads to inhibition of the propagation of prion seeding (11, 52, 53). To test the effect of guanidine, we first empirically determined the optimum concentration of guanidine. We found that while 5mM GnHCl reduced cell viability of *S. pombe*, cells grew normally at 4mM GnHCl (not shown). Therefore, cells of the parental strain SPA236, two independent [SMG2]M segregants from tetrads generated from the cross, asshown in **Figure 3B**, two independent [SNG2]° derivatives generated by *cut4* overexpression in cells of genotype I, followed by plasmid loss (**Figure 4A, 4B**) and two independent colonies of a WT prototrophic ura^+^ strain were grown in the presence of 4mM guanidine and effect on spo^+^ and ura^+^ phenotypes was analyzed by plating the treated cells on appropriate medium. The parent strain SPA236 having low level of spo^+^-ura^+^ (5-10% of colonies) exhibited no change in the level of spo^+^-ura^+^ phenotypes on guanidine hydrochloride (**Figure 5A-5C**). Likewise, as controls, the WT ura^+^, h^90^ strains (which sporulate normally and form spo^+^ colonies, showed similarly high levels of ura^+^ phenotype and low percentage of colonies with spo^-^ phenotype before and after the guanidine treatment (**Figure 5B, 5C**). Interestingly, the putative [SNG2]_M_ derivatives showed reduced growth on plates lacking uracil and loss of the spo^+^ phenotype following guanidine treatment; a similar effect was observed in case of [SNG2]° cells (**Figure 5A-5D**), which were derived from [SNG2]_M_ cells generated by overexpression of *cut4* gene, followed by plasmid curing (**Figure 4C**). However, cells of DSPR showed no effect of guanidine treatment on their spo^+^ and ura^+^ phenotypes (**Figure 5A-5C**).

**Figure 5.**
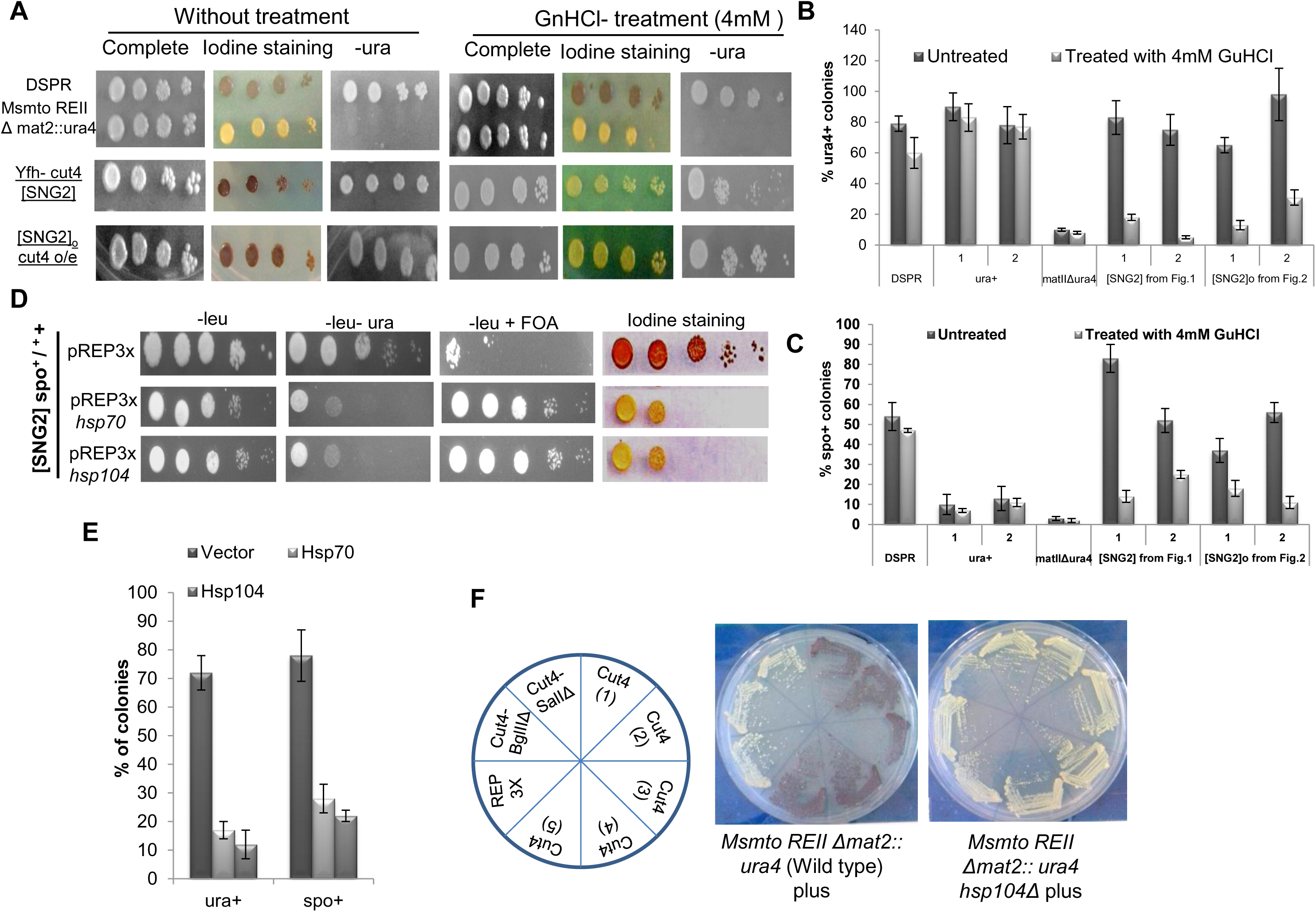
Curing of [SNG2] prion by Gu-HCl and overexpression of heat shock proteins. **(A)** Serial dilution spotting assay of DSPR, *Msmto REIIΔ mat2P::ura4*, SNG2]M-YFH-cut4 (spo^+^-ura^+^) segregants and [SNG2]° (*cut4* o/e) along with the parental strains on complete plates and plates lacking uracil, with and without treatment of 4mM Gu-HCl. o/e denotes overexpression. **(B)** Quantitation of percentage of colonies with ura^+^ phenotype on treatment of various strains with 4mM Gu-HCl. ura^+^ represents control WT, DSPR, SPA236 with genotype *Msmto ura4D18 REIIΔ mat2P::ura4*, and two independent [SNG2]_M_ (spo^+^-ura^+^) segregants from **Fig. 2** and two independent [SNG2]^0^-GFP-cut4 from transformants showing ura^+^ phenotype after plasmid loss, as shown in **Fig. 4D**. **(C)** Quantitation of percentage of spo^+^ colonies on treatment of different indicated strains with 4mM Gu-HCl. (Iodine staining indicates staining of colonies with iodine after 3 days of growth). **(B, C)** Data was represented as SD. **(D)** Curing of prion form of [SNG2]_M_ by overexpression of *hsp* genes in segregant 3B (spo^+^-ura^+^) obtained from cross shown in Figure 2 assessed by serial dilution spotting assay. **(E)** Quantitation of the effect of overexpression of hsp70 and hsp104 in [SNG2]_M_ strain on percentage of colonies showing spo^+^ and ura^+^ phenotype. Error bars represent SD. **(F)** Iodine staining of transformants generated from expression of pREP3X, *cut4 DB/B* and *cut4* in WT strain with genotype I (*Msmto REIIΔmat2P::ura4*) and same strain having *hsp104Δ* mutation (*Msmto REIIΔmat2P::ura4 hsp104Δ)*. Independent transformants were streaked on selective plates (PMA-leu) and colonies stained after 4 days’ growth at 30°C.

### Curing of [SNG2] prion-form by overexpression of Hsp70 and Hsp104

Overexpression of heat shock proteins Hsp70 and Hsp104 has been shown to cure the prion phenotypes and promote the solubilization of the prion aggregates (21, 49-54). Therefore, we transformed the [SNG2]_M_ strain, having spo^+^-ura^+^ phenotype, with empty vector and *hsp104* and *hsp70* genes of *S. pombe* and *S. cerevisiae,* respectively. We monitored the phenotypes of the transformants and the aggregation pattern of prion protein. Consistent with the reported effect of overexpression of *hsp* genes on curing of the prion structure (49) there was a 6-8 fold reduction in the number of transformants having spo^+^-ura^+^ phenotype similar to the background level observed in the WT parent strain SPA236 (**Figure 5E**).

It has been shown that generation and propagation of prions requires presence of molecular chaperones, such as heat shock proteins (55) as they are known to shear the aggregates into small fibrils for transmittance to filial generation. We compared the generation of spo^+^ phenotype by overexpression of *cut4* gene in the wt strain having genotype I (SPA236) to the same strain having *hsp104Δ* mutation. Results show that while all the transformants of the strain SPA236 give spo^+^ phenotype, those having *hsp104Δ* mutation did not (**Figure 5F**). Thus, the generation of [SNG2]_M_ by overexpression of *cut4* is dependent on *hsp104*.

### Formation of Amyloid aggregates and their solubilization by Hsp104

Several studies have shown that the prion forms of proteins adopt amyloid structure showing retarded electrophoretic mobility under semi-denaturing conditions in agarose gels (Semi denaturing detergent agarose gel electrophoresis, SDDAGE), where in the samples are mixed with 1% SDS but, unlike SDS-PAGE, not boiled prior to electrophoresis in the presence of 0.1% SDS. Unlike earlier reports, we carried out SDDAGE in presence of 2% sarkosyl instead of 1% SDS, as we failed to observe aggregates under the latter conditions. Results showed that while the parent strain having YFP-tagged Cut4 from the cross shown in Figure 4A showed just a broad band but no aggregates (**Figure 6A**, lane 1), the segregants 2A, 2B, 3B and 4B, with the phenotypes ura^+^-spo^-^, ura^+^ spo^+^, ura^+^-spo^+^ and ura^w^-spo^+^, respectively, all showed a broad band of aggregates (**Figure 6A**, lanes 2-5). No band was observed in the untagged parent strain shown in Figure 4A (**Figure 6A**, lane 6). It is interesting that the segregants showing weaker phenotype (**Figure 6A**, segregants 2A and 4B) have lower intensity of the aggregate band as compared to the segregants with stronger phenotype (**Figure 6A**, segregants 3B and 4B). Surprisingly, in addition to high molecular weight region, the aggregate bands also spans the monomer and sub-monomer region.

**Figure 6.**
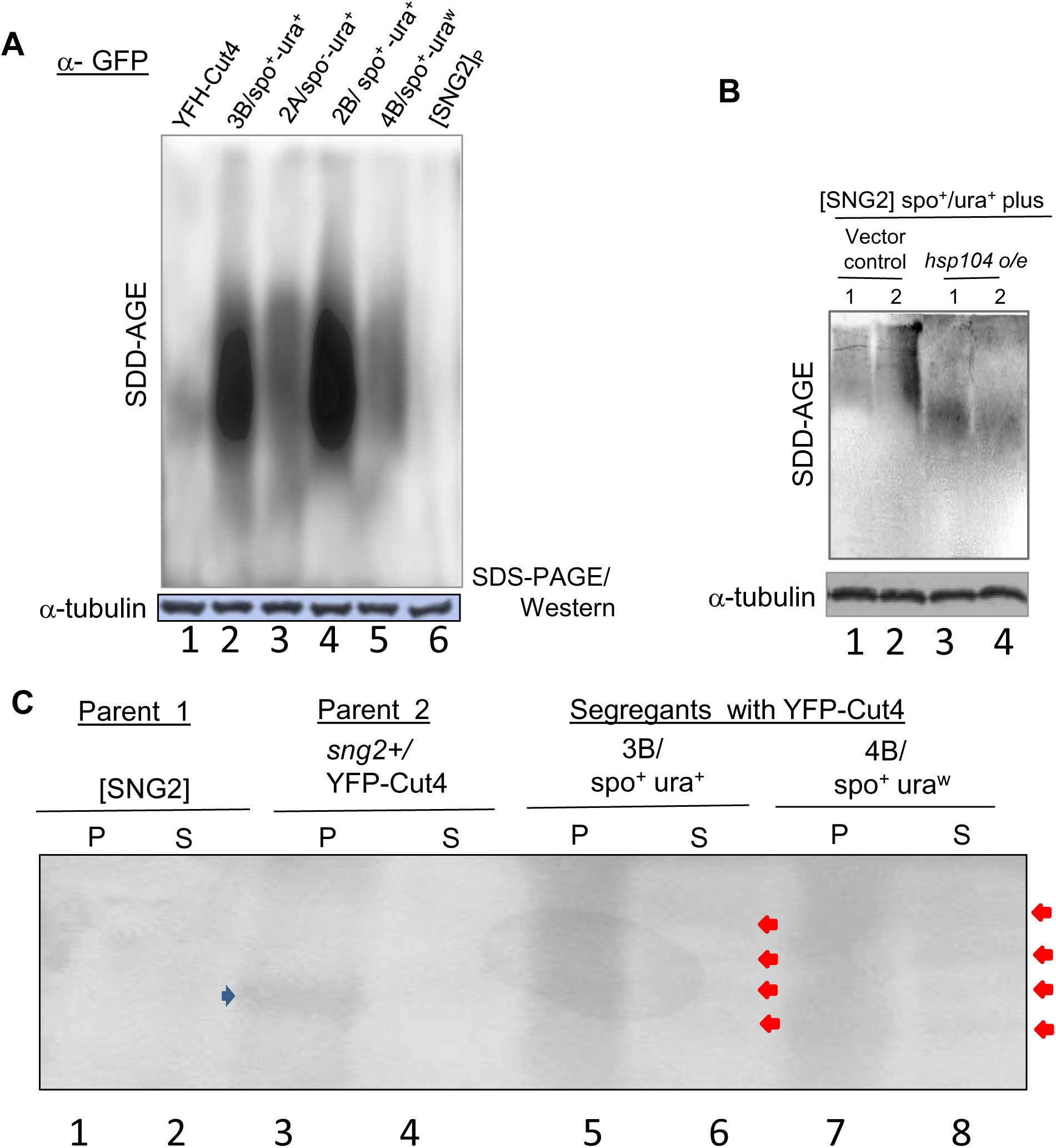
Formation of Cut4 amyloid aggregates in [SNG2] prion form. **(A)** SDD-AGE followed by western blotting with anti-GFP antibody of the protein extracts from segregants shown in Figure 4B. Parent 1: sng2^+^/ YFH-Cut4 (lane 1), segregant 3B, showing spo^+^-ura^+^ phenotype (lane 2), segregant 2A showing spo^-^-ura^+^ phenotype (lane 3), segregant 2B showing spo^+^-ura^+^ phenotype (lane 4), segregant 4B showing spo^+^-ura^w^ phenotype (lane 5) and parent 2: [SNG2]_P_ lacking YFP-tagged Cut4 (lane 6). **(B)** Conversion of amyloid aggregates to monomeric form by overexpression of *hsp104*. SDDAGE analysis of protein extracts of the two independent [SNG2]_M_ strains harbouring GFP-Cut4 from Figure 5D, transformed with empty vector (lanes 1 and 2) and vector overexpressing *hsp104* (lanes 3 and 4). **(C)** SDDAGE analysis of soluble and pellet fraction from the parent strains and segregants of the cross shown in Figure 4A and 4B). Arrowheads indicate the oligomeric forms of YFP-Cut4 in the soluble fraction.

Furthermore, it has been shown that heat shock proteins hsp70 and hsp104 can convert the amyloid aggregates into monomers (49). We subjected two independent ura^+^-spo^+^ transformants obtained by overexpression of *cut4* in strain having GFP-tagged *cut4* gene (**Figure 4C, 4D**) to loss of the *cut4* plasmid and retransformation with empty vector or *hsp104* gene. SDDAGE analysis followed by western blotting with anti-GFP antibody showed the presence of aggregates, which spanned the region across the gel up to the sample well in strains transformed with the empty vector (**Figure 6B**, lanes 1 and 2). Interestingly, monomer sized bands were observed in strain transformed with *hsp104* gene (**Figure 6B**, lanes 3 and 4).

The apparent discrepancy between the electrophoretic mobility of the aggregate bands in Figure 6A and 6B can be explained by the shorter duration of the electrophoresis in the latter; the longer duration of electrophoresis in Fig. 6A could have led to splitting and diffusion of the smaller oligomers of Cut4 through the agarose gel, which did not occur due to the shorter run length of electrophoresis in Figure 6B.

We further carried out fractionation of cell extracts of the parent strains and the prionic segregants from Figure 4A into soluble and pellet fractions followed by SDDAGE and western blotting with anti-GFP antibody. In the parent strain YFP-Cut4 was found entirely in the pellet fraction. Interestingly, among the prionic segregants 3B and 4B (**Figure 4A**), while the pellet fraction contained only broad aggregate band (**Figure 6C**, lanes 5 and 7), sharp bands were detected in the soluble fraction (arrowheads, **Figure 6C**, lanes 6 and 8). The sharp bands may represent distinct soluble oligomeric intermediates, while the broad aggregate band may represent a polydisperse spreading of YFP-Cut4 aggregates.

### Dominance and cytoplasmic inheritance of [SNG2] - Cytoduction Experiment

Because of their cytoplasmic mode of segregation, the prion forms show dominance and cytoplasmic inheritance (9, 15, 17). This has been demonstrated by cytoduction experiment using a karyogamy defective mutant *kar1* in *S. cerevisiae* (3, 11). It has been shown that, similar to the *kar1* mutant of *S. cerevisiae*, the *tht1Δ* mutant is defective in karyogamy in *S. pombe* (56), as it shows lack of fusion of the parental nuclei and meiosis without crossing over. We confirmed this characteristic of the *tht1Δ* mutant and observed a lack of intrachromosomal recombination between *leu1* and *his2* markers on *chrII* and parental co-segregation of *chrIII* (*ade6)* and *chrII* marker (*leu1^-^* and *his2^-^*) in a meiotic cross (**Figure S4**).

Stable diploid strains homozygous for *tht1Δ* and genotype I (*leu1-32 mat1Msmto REIIΔmat2P::ura4*) were constructed by cell fusion and interallelic complementation of the *ade6-210* and *ade6-216* alleles (34), wherein the dominance and cytoplasmic segregation of silencing defect caused by [SNG2] and *swi6Δ* mutation was investigated in heterozygous condition. While a dominant phenotype would manifest as spo^+^ (due to meiosis in the azygotic diploids; see materials and methods) and ura^+^ phenotype, like that in haploid mutant, the spo^-^-ura^-^ minus phenotype would imply recessiveness of the mutation (**Figure 7A**). Interestingly, the heterozygous diploid [SNG2]_M_/*cut4*^+^ shows spo^+^-ura^+^ phenotype, indicating the dominance of the [SNG2] over wt cut4. On the other hand, like the parent diploid strain, the *swi6Δ*/*swi6^+^* diploid shows spo^-^-ura^-^ phenotype, indicating that the *swi6Δ* mutation is recessive (**Figure 7B, 7C**). Microscopic examination showed the occurrence of azygotic asci in spo^+^ colonies of *tht1Δ*/*tht1Δ*, [*SNG2*]M/*cut4^+^* diploids, which implies that defect in silencing of *mat2P* locus in [SNG2]_M_ is dominant over *cut4^+^* (**Figure 7B**). As expected, the *[SNG2]* haploid and *swi6Δ* haploid strains show iodine staining due to defective silencing (**Figure 7B**, upper panel), while the [SNG2]_P_/P, *tht1Δ/tht1Δ* did not does not show iodine staining and loss of silencing since both *mat1* and *mat2* harbor the same mating type information Plus (**Figure 7B**, lower panel). **Figure 7B, lower panel)**. These results support the cytoplasmic inheritance and dominance of the [SNG2]_M_, while confirming the recessive nature of the *swi6Δ* mutation.

**Figure 7.**
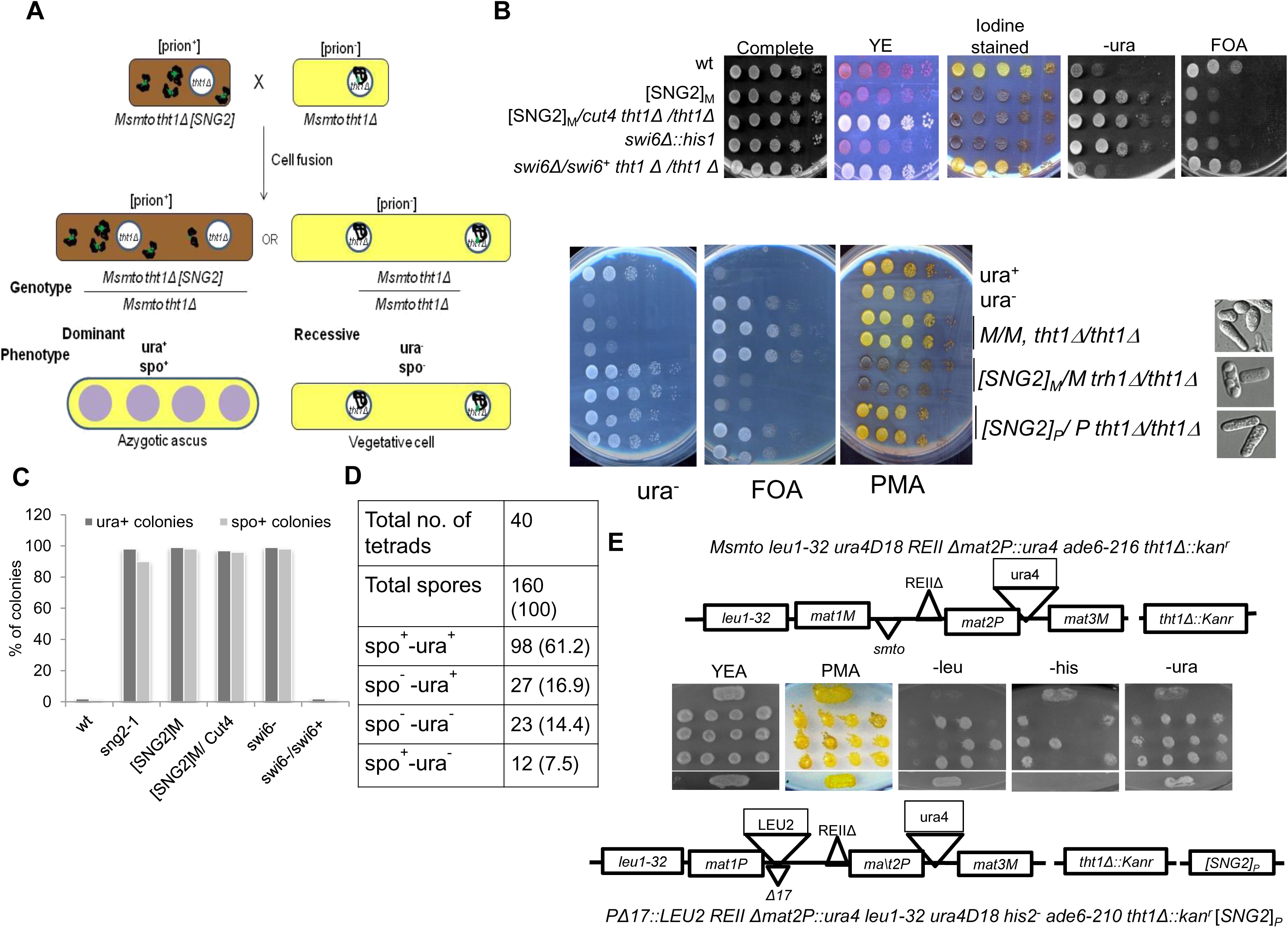
Cytoplasmic inheritance and dominance of [SNG2]. **(A)** Diagrammatic representation of cytoduction experiment for generation of dominant (prion^+^ form) and recessive (prion^-^ form) in karyogamy deficient (*tht1Δ*) diploid strain. **(B)** Serial dilution spotting assay of strains with indicated genotypes on complete, YE, PMA^+^, -ura and FOA plates. Colonies growing on PMA^+^ plates were stained with iodine to check sporulation. Lower panel includes the control strains. Pictures on the right side and photomicrographs of the cells subjected to sporulation conditions to induce meiosis. **(C)** Quantitation of ura^+^ and spo^+^ colonies of strains with indicated genotypes after growth on plates lacking uracil and sporulating plates, respectively. **(D)** Tetrads derived from a cross of [SNG2]_P_-*tht1Δ* strain (prion form with genotype II having *tht1Δ*) having spo^-^-ura^+^ phenotype and a WT strain with *tht1Δ* (genotype I) having spo^-^-ura^-^ phenotype, were spotted on YEA and then replica plated to different plates, as indicated. **(E)** Segregation pattern of ura^+^ phenotype among *tht1Δ* mutants in tetrad analysis represented in **(D)**.

A further test for the cytoplasmic mode of inheritance of [SNG2]_M_ was performed by setting up tetrad analysis of a meiotic cross between a WT *tht1*Δ strain with genotype I and a *tht1*Δ, [SNG2]_P_ strain. Very interestingly, as shown in the normal cross (**Figure 3B**), several spo^+^-ura^+^ segregants were obtained (**Figure 7D, 7E**) with ∼68.7% of segregants being spo^+^ and 78% segregants showing the ura^+^ phenotype (**Figure 7D**) and non-Mendelian segregation (**Figure 7D, 7E**). These results further support the dominance and cytoplasmic basis of inheritance of the [SNG2] prion form.

As another test for cytoplasmic inheritance as well as to test whether the prion form could confer silencing defect at other heterochromatin loci, like *mat3* locus (*mat3::ade6*) and *cen* locus (*otr1R::ade6*), we first crossed cells of [SNG2]_P_ with a naïve WT strain having Minus mating type (*mat1Msmto*) without any reporter gene. Five independent derivatives having *mat1Msmto* locus and no reporter from the above cross, tentatively labeled [SNG2]_M_ 1-5, were then crossed with strains having Plus mating type along with either the *otr1R::ade6* locus (57; **Figure S5C, panel b)** or *mat3::ade6* locus (58; **Figure S5C, panel a)**. Random spores from these mating as well as the parental strain with the *mat3::ade6* or *otr1R::ade6* locus were streaked on YE plates to assess the effect on silencing of the *ade6* reporter at *otr1R* (**Figure S5A**) *and mat3* (**Figure S5B**) locus, respectively. In both cases, while the parental strains give dark red colonies indicating a transcriptionally silent *ade6* reporter (labeled control, **Figure S5A, S5B**), the cross involving the derivatives [SNG2]_M_1-5 produced progeny with variable degree of white and pink colonies representing varying levels of derepression of the *ade6* reporter both at *mat3* and *otr1R* loci **(Figure S5A-S5C)**.

To further confirm that the [SNG2] prion can cause silencing defect at the centromere locus, we overexpressed *cut4* gene in the strain having *otr1R::ade6* reporter. A high rate of generation of pink colonies in transformants with high copy vector containing *cut4* gene, but not one containing truncated *cut4 DB/B* gene or the vector control (not shown) in the strain SPA236, as shown earlier, confirms that the putative prion form generated upon overexpression of *cut4* gene can trigger loss of silencing at the centromere locus as well.

### Propagation of [SNG2] by protein infection

A key test of protein-based inheritance is the dominant ability of protein obtained from the [prion^+^] cells to impart the phenotype upon transformation into the naïve [prion^-^] cells (59). In the present experimental set up, ability of protein extract from [SNG2]_M_ or [SNG2]° cells to induce the phenotype of spo^+^, associated with haploid meiosis and ura^+^ phenotype when transformed into strain with genotype I (*leu1^-^ mat1Msmto REIIΔmat2P::ura4*), would indicate its infectivity, while inability to do so would indicate it to be non-prion form (Schematic representation, **Figure 8A**). Accordingly, we co-transformed cells of strain SPA236 (genotype I) having GFP-tagged *cut4* gene, with empty vector pREP3 along with benzonase-treated protein extracts prepared from cells of WT, [SNG2]_P_, [SNG2]°, DSPR, LSPR and the [SNG2]_M_ segregants obtained from meiotic cross shown in **Figure 2**. The selected leu^+^ transformants were screened for the phenotype indicative of silencing. Results show that while cell extracts from WT cells had no effect on expression of the *ura4* reporter and on iodine staining phenotype, extracts of cells of [SNG2]_P_, [SNG2]°, [SNG2]_M_, all caused elevated level of spo^+^-ura^+^ colonies, as indicated by iodine staining, enhanced growth on plates lacking uracil or lack of growth on FOA plates (**Figure 8B**).

**Figure 8.**
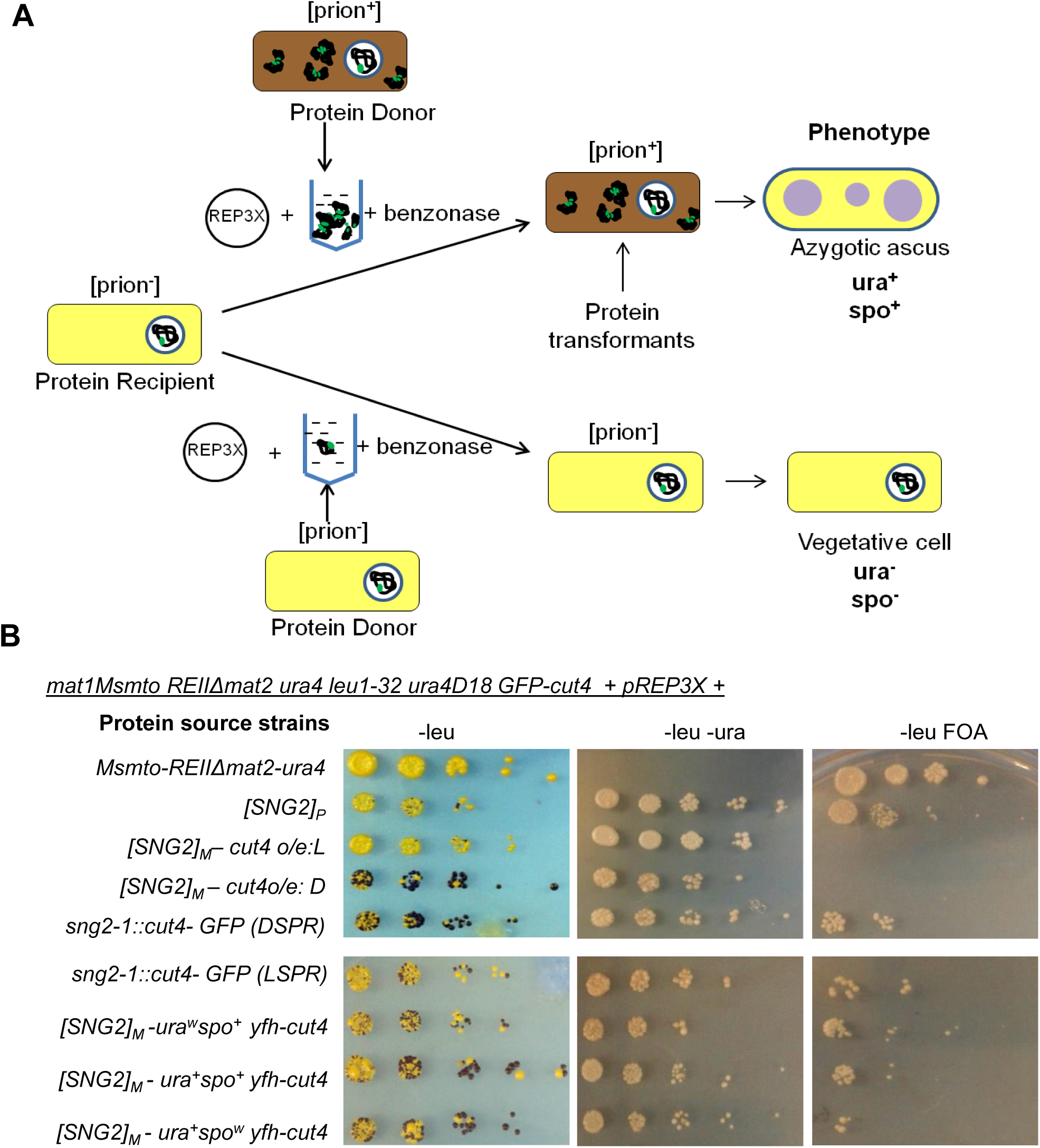
Propagation of [SNG2] prion by protein transformation. **(A)** Diagrammatic representation of test for infectivity of [SNG2] prion, as described in Materials and Methods and Results. **(B)** Serial dilution spotting assay of wild type strain (genotype I with GFP tagged *cut4*) transformed with protein from indicated strains, on complete PMA^+^, -ura and FOA plates. Colonies growing on PMA^+^ minus leu plates were stained with iodine to check sporulation.

Notably, even extract from cells of DSPR induced the spo^+^-ura^+^ phenotype strongly, while that from cells of LSPR was less effective (**Figure 8B**). This result suggested that the Dark, spo^+^-ura^+^, derivatives of *sng2-1* mutant with genotype I contained the prion form by itself, while the LSPR may have a lower fraction of prion-form as it only exerts a modest effect. The intermediate level of growth on plates lacking uracil and spo^+^ phenotype on protein transformation with LSPR protein extracts mimics the behavior of the parent LSPR strain; **Figure 1D**). Thus, the two epigenetic states of the *sng2-1* mutant, DSPR and LSPR, initially observed by us (**Figure 1**) may predominantly represent the strong and weaker prion forms, respectively, of the mutant Cut4p.

### [SNG2] prion-form phenocopies *cut4* mutation and affects stress tolerance

In general, the prion forms are associated with loss of function of the WT protein function. As *cut4-533* mutant has been associated with ‘*cut*’ (‘chromosomes untimely torn) phenotype and enhanced cadmium sensitivity, we checked whether the cells of the [SNG2] prion form may exhibit similar phenotypes. Indeed, we find that cells of both [SNG2]_P_ and [SNG2]_M_ exhibit enhanced ‘*cut*’ phenotype (**Figure 9A**) as well as sensitivity to Cd^2+^ (**Figure 9B**) as compared to the WT parent and closely phenocopies the *cut4-533* mutant (29). In addition, they also show enhanced resistance to oxidative, thermal and ethanol stresses (**Figure 9C-9E**). Furthermore, like the heterochromatin mutant *swi6^-^*, the [SNG2] cells also show enhanced rate of chromosome loss (**Figure 9F**). Thus, the prion-form [SNG2] mimics a functionally defective APC and heterochromatin mutant.

**Figure 9.**
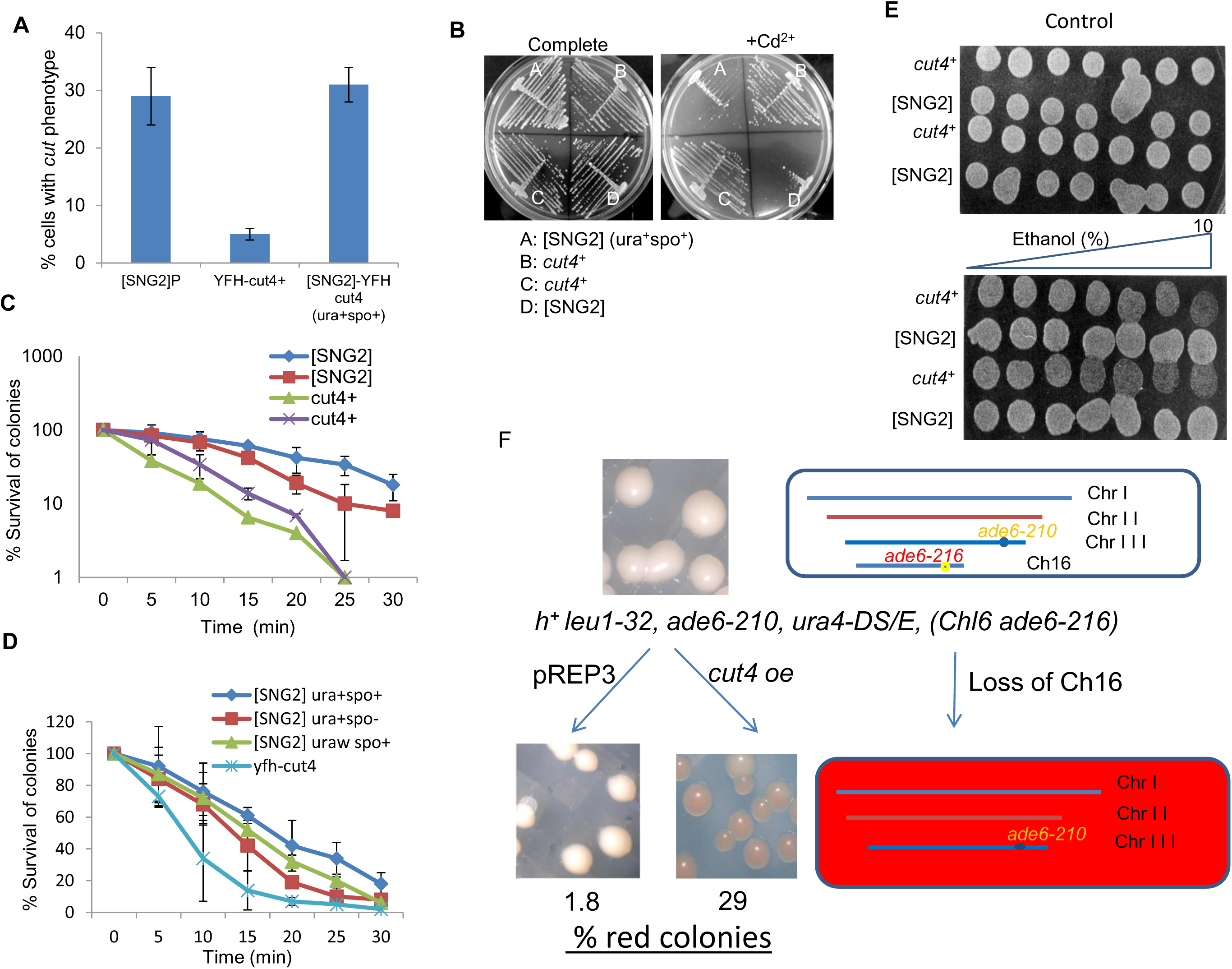
Cells of [SNG2] prion form phenocopy *cut4* mutant and show enhanced stress response. **(A)** Histogram representing the percentage of cells having ‘cut’ phenotype in indicated strains. **(B)** Growth of WT and [SNG2] segregants of a single tetrad on PMA plate containing 20μM CdCl_2_. **(C)** Plot representing survival of WT and [SNG2] cells subjected to heat stress at 52°C for the indicated time periods. **(D)** Plot representing survival of WT and independent [SNG2] segregants upon exposure to 1mM H_2_O_2_ for the indicated time periods. **(E)** Spotting assay of cells of two WT and [SNG2] strains each on control plate (top panel) and plate containing 0-10% gradient of ethanol (Lower panel). **(F)** Enhanced rate of chromosome loss in [SNG2] cells. The schematic depicts the basis of screen for chromosome loss. Cells of mutant showing enhanced loss of the artificial chromosome (Ch16 ade6-216) form red colonies, while WT cells with high level of chromosome stability produced mostly white colonies on the adenine-limiting YE plate (75).

## DISCUSSION

Following up on t h e discovery of t h e copper transport associated protein Ctr4 as the first prion in *S. pombe* (60), here we report [SNG2] as a prion-form of Cut4 protein from *S. pombe,* which abrogates heterochromatin silencing and affects stress tolerance similar to the *sng2-1*/*cut4* mutant. Earlier, Sideri *et al*. (60) carried out a predictive analysis and provided a list of probable prion protein (on the basis of Q/N rich prion domains), which does not include Cut4p as a probable prion protein. However, prions [Het-s] from Podospora and PrP^Sc^ from human also lack Q/N rich sequences but are rich in highly charged residues. Likewise, Cut4p is also lacking in Q and N residues but is rich in β- strands (not shown).

Nonetheless, we have shown that the *sng2-1* mutant of *cut4* is a variant that efficiently produces the [SNG2] prion. We have also demonstrated that distinct variants of [SNG2] prion exhibit varying phenotypes and aggregation levels of Cut4p. The main significance of this study is that in addition to the chromosomal basis of inheritance, silencing and heterochromatin structure can also be regulated cytoplasmically in a prion-like manner by [SNG2], the prion form of Cut4p, in *S. pombe*.

The findings in the present study originate from the isolation of *sng2-1* a ts mutant allele of *cut4*, which exhibited two states of silencing: the derepressed (DSPR) having infectious aggregates of [SNG2] and repressed (LSPR) with less aggregated and less infectious Cut4^sng2-1^ protein state. The two variants DSPR and LSPR show distinct phenotypes; spo^+^-ura^+^ and spo^-^-ura^w^, respectively, representing strong sporulation (spo^+^) and *ura4* expression (ura^+^), and lack of sporulation (spo^-^) and weak ura expression (ura^w^), as reflected in the level of iodine staining and growth on medium lacking uracil. The dark spo^+^-ura^+^ state, denoted here as DSPR, displayed dominance, as it could only be partially complemented by the *cut4* gene on integrating vector but not on high copy vector. Importantly, the spo^+^-ura^+^ phenotype persisted even after the ts^-^ phenotype was segregated out, suggesting that it may have generated a self-propagating prion state. A similar scenario occurs upon overexpression of *cut4* gene on a high copy plasmid.

Thus, the DSPR probably represents a prion precursor form of the mutant protein, which converts the normal Cut4 into prion form. The purified derivatives obtained by crossing and having spo^+^-ura^+^ phenotype were found to fulfill the following prion-like characteristics i) Non-Mendelian inheritance, ii) dominance and cytoplasmic inheritance, iii) curing by guanidine and overexpression of *hsp70* and *hsp104* genes and dependence of propagation of prion-form on *hsp104*, iv) generation of prion form by overexpression of *cut4* gene, v) aggregate formation by the prion form of Cut4 and vi) regeneration of the phenotype by protein transformation. These results lend support to the role of Cut4 as a prion. The non-Mendelian inheritance is not complete, similar to that reported for [URE3] prion by Wickner (11).

Earlier studies in fission yeast showed that heterochromatin structure is regulated in *cis*. For example, Grewal and Klar showed that a strain carrying a deletion of the region spanning the *mat2*-*mat3* interval exhibited two alternative states of chromatin that exhibit repressed and derepressed phenotypes (22). Most interestingly, these states were stably propagated during mitosis and meiosis. Furthermore, the role of RNAi at the level of establishment of the repressive epigenetic mark of H3-Lys9 is also shown to be chromosome borne (25). Parallel to RNAi, the histone deacetylase Clr3 is recruited by the ATF/CREB family proteins and, thereafter, extends in *cis* along with Swi6/HP1 to establish heterochromatin (61).

Earlier work from this lab showed that APC subunits Cut4 and Cut9 interact with Swi6 and the APC and Swi6 act in a mutually cooperative manner to establish heterochromatin structure at the centromere and mating type loci (30). The present study shows that a prion form of Cut4 can lead to loss of silencing. It may be speculated that lack of interaction of the prion from of Cut4 with Swi6 may lead to failure to recruit Swi6 to heterochromatin, resulting in loss of silencing. It is also likely that [SNG2], the prion form of Cut4, is defective in the assembly of APC, as the [SNG2] cells closely phenocopy the *cut4-533* and *sng2-1* mutants. Further investigation will be needed to test these possibilities.

### Variability of [SNG2] prion phenotypes

Surprisingly, we observe considerable variation in phenotypes of the [SNG2] derivatives-at least 3 types of phenotypes were observed among the meiotic progeny of the cross of [SNG2]_M_ with SPA302 (genotype II), or [SNG2]_P_ with SPA236 (genotype I): spo^+^-ura^+^, spo^+^-ura^w^ and spo^-^-ura^+^. The spo^+^-ura^+^ phenotype represents derepression of both *mat2Pc* and the *mat2P*-linked *ura4* reporter, with the spo^+^- ura^w^ representing derepression of *mat2Pc* but weaker derepression of the *ura4* reporter, while spo^-^-ura^+^ represents the derepression of *ura4* reporter but not *mat2Pc*. Such diversity in prion phenotypes has been observed in other prions (15, 62). In case of [SNG2], this variation may represent varying extent of heterochromation assembly defect. Interestingly, we find that the specific expression states (eg. spo^+^-ura^+^, spo^+^- ura^w^, spo^-^-ura^+^) can be stably inherited in genetic crosses indicating different stable and heritable form of the [SNG2] prion with distinct structure and function (data not shown).

As heterochromatin assembly involves establishment followed by expansion through components like *clr3*, *clr4*, *swi6* etc. Heterochromatin assembly at mating-type region involves action of at least four pathways: (i) *mat2*- linked (31, 33) (ii) RNAi- mediated action at the *cenH* region (28, 63), (iii) ATF- CREB pathway acting at *mat3* (64, 65) and (iv) Pap1-dependent pathway (66). Thus, the variability of [SNG2] effect at the *mat3*-linked *ade6* reporter could imply different degrees of effect on expansion of heterochromatin across the *mat2-mat3* interval. Furthermore, the [SNG2] also acts in a similar manner to derepress the *cen::ade6* and *mat3::ade6* reporters, and thereby the heterochromatin structure at the *cen1* and *mat3*, respectively. However, the transmission efficiency of the spo^+^-ura^+^ phenotypes during meiotic cross ranged from 62-86% and during protein infectivity assay was around 46-71% among independent transformations, although they originate from crosses between closely related strains, ruling out the possibility of yeast viruses mediating the transmission which is reported to be 100% (67).

It is worth noting that while the native WT YFP-Cut4 was fractionated almost entirely in the pellet fraction (**Figure 6C**, compare lanes 3 and 4), the protein from the spo^+^-ura^+^ segregants showed partitioning as broad aggregates in the pellet fraction (**Figure 6C**, lanes 5 and 7) but also as distinct bands in the soluble fraction (**Figure 6C**, lanes 6 and 8). This is consistent with the observed high prionogenicity of the smaller oligomer derivatives (46).

### Presence of Intrinsically disordered sequences in Cut4p

The inheritance of prion proteins requires both the generation and propagation of the amyloidogenic aggregates. Many of proteins form aggregates but they do not propagate in a non-Mendelian fashion to their filial generations. These include inclusion bodies, folding aggregates and amorphous bodies. Another class of proteins, the intrinsically disordered proteins can also form aggregates, which propagate in a non-Mendelian fashion to their daughter cells and their propagation is dependent on molecular chaperone, some of which can form amyloids (21, 68).

We have investigated the presence of aggregate-forming and amyloidogenic domains in Cut4p, utilizing various bioinformatics tools. We find that Cut4p lacks long Q/N stretches (60) that are associated with some prion forming proteins. Use of the PAPA algorithm of PLAAC software, which is based on detection of Q/N rich regions, yielded a moderate prediction score of 0.051. Conventionally, segments with score of above 0.05 have high propensity to form PrLDs **(Supplementary Figure S7**). However, PLACC also provides prediction from other algorithms, like Foldindex, which predicts intrinsically disordered regions (IDRs). Thus, the regions underlined in green are prion-like based on domain prediction from PAPA, while regions underlined in black are predicted as IDR regions by Foldindex **(Supplementary Figure S7**), as also by the DISOPRED software. Interestingly, IDRs have also been associated with potential to form prions (69).

Analysis using various aggregate and amyloidogenic region predictor tools such as Amylopred2, Tango, Fold Amyloid, Zygoggregator,etc. also reveal that Cut4p of *S. pombe* contains similar stretches with high amyloidogenic potential (results not shown). Thus, Cut4p contains two regions – residues 2-395 and 1251-1380 that are rich in β-strands. In future studies, we will investigate the role of the domains identified by Foldindex and DISOPRED software in inducing prions by doing domain swapping experiments with SUP35.

Curiously, the *sng2-1* mutation at residue 984 does not map to any one of the regions predicted above. We suggest that the *sng2-1* mutation may accentuate the potential of the putative prionogenic regions to form prions, as shown earlier (70). Furthermore, it is interesting that the *sng2-1* mutation (amino acid residue 984) is located in the region spanned by the *Bgl*II sites, whose deletion abolishes the prionogenic potential of Cut4, as shown in Figure 4.

### Cellular Consequences of prion form of Cut4

It is a moot question whether the aggregated prion form of Cut4 performs its normal physiological functions. We find that the [SNG2] cells display similar sensitivity to heavy metal ions like, Cd^2+^, as the *sng2-1* and *cut4-533* mutants (29, 71).

Thus, the prion form of Cut4 phenocopies the *cut4* mutant representing a loss of function, as expected in case of prions. Likewise, similar to the *cut4* and other *‘cut’* mutants, the [SNG2] cells also show high incidence of‘*cut*’ (chromosomes untimely torn) phenotype, which represents deregulation of cell cycle progression (29, 71). Furthermore, like *sng2-1* and *swi6* mutants, [SNG2] cells also show enhanced rate of chromosome loss indicating that the prion-form of Cut4 causes chromosome instability, like silencing mutant *swi6* (30).

### Evolutionary significance

It has been proposed that some prions confer advantage on cells to survive and adapt to various stresses (72). In agreement, we find that cells of the [SNG2] prion exhibited enhanced tolerance to various environmental stresses, like ethanol stress, thermal stress and oxidative stress. It is puzzling that Cut4, a protein performing an important and evolutionarily conserved role of regulating cell cycle progression in all eukaryotes, has evolved to retain regions that makes it apparently non-functional and imparts stress tolerance. Computational analyses reveals that regions having potential to form amyloids are conserved in Cut4 sequences in other species as well (not shown), thus supporting the notion that such sequences have been selected during evolution and may provide selective advantage in response to environmental changes.

Paradoxically, the [SNG2] cells also exhibited cell cycle defect and enhanced chromosomal loss, conditions characteristic of disease states like cancer. It is possible that cell cycle defects and chromosomal defects may confer enhanced stress tolerance, a trait exhibited by cancer cells. The enhanced stress tolerance of cells bearing [SNG2] suggests that the [SNG2] prion may facilitate the adaptation of the *S. pombe* cells to various environmental changes, thus providing a basis for selection of their propagation during mitosis and meiosis.

That Cut4^sng2-1^ mutant protein may adopt a metastable prion structure more efficiently than WT Cut4p is not unprecedented. For example [PSI^+^] has been shown to arise 5000-fold more frequently in the Sup35p mutant containing two additional oligopeptide repeats (73). Likewise, H2p mutant of Ure2p causes 1000-fold increase in the rate of [URE3] induction although this and other similar mutations lie outside the prion domain (70). It has been suggested that conformational flexibility may enhance the frequency of prion induction.

In conclusion, this is the first report of a prion affecting heterochromatin structure in eukaryotes. The ease of doing genetics, biochemistry and cell biology in fission yeast should facilitate a deeper insight of molecular mechanisms and significance of Cut4p prion both in yeast in future studies.

## ACKNOWLEDGEMENTS

We thank M. Yanagida and D. Masison for strains and plasmids. We are grateful to Liming Li and Deepak Sharma for helpful suggestions and Dr. Amar Klar for critically reading the manuscript. RND and PS were recipients of SRF from CSIR and SS for RA fellowship from DBT. Harish Kumar and Dr. Ram Yadav are acknowledged for help with confocal microscopy. Dr. Gurprit is acknowledged for help in bioinformatic analysis.

## Author contributions

S.S. performed experiments, analyzed data, prepared the figures, wrote the paper; R.N.D carried out experiments and analyzed data, P.S. performed the experiments and analyzed data and J.S. designed the research, analyzed data, and supervised the work.

## Competing interests

The authors declare no competing financial interests.

## Notes

### Competing Interest Statement

The authors have declared no competing interest.

