## Supplementary file for "[SNG2], a prion-form of Cut4/ Apc1, enforces non-Mendelian inheritance of heterochromatin silencing defect in Fission Yeast"

### Supplementary Data

#### High mitotic stability of the [SNG2] prion forms

We determined the stability of the $[SNG2]_M$ prion. We found that three independent [SNG2] segregants from the cross shown in **Figure 2** showed stable though variable phenotypes, with 66-98% and 60-96% of colonies showing $spo^+$ and $ura^+$ phenotypes, respectively, over repeated freeze- thawing cycles (not shown). The original forms of the *sng2-1* mutant (DSPR and LSPR) and various [SNG2] derivatives with $spo^+$-$ura^+$, $spo^-$-$ura^+$, $spo^+$-$ura^w$ and $spo^w$- $ura^+$ phenotypes were inherited stably during mitosis, showing a switch to the opposite state at rate/generation ranging from $1.7X\ 10^{-4}$ to $5.6X10^{-5}$/generation for $spo^+$ phenotype and $1X10^{-4}$ to $9.5X10^{-5}$/generation for $ura^+$ phenotype, respectively **(Figure S1B)**. Similarly, the [SNG2] prion phenotype was also stably propagated during meiosis: nearly 56% and 23% segregants of a meiotic cross retained the $spo^+$ and $ura^+$ phenotypes, respectively, during meiosis. Similar stable pattern of inheritance was observed during repeated crosses.

**Table S1. Strains used in the study**

| Strain | Genotype |
|---|---|
| **SPA 236** | *mat1Msmto leu1-32 ura4D18 REII Δmat2P::ura4 ade6-216* |
| **SPA302** | *mat1PΔ17::leu2 REIIΔmatP2::ura4 leu1-32 ura4D18* $his2^-$ *ade6-210* |
| **FY566** | $h^+$ *leu1-32, ade6-210, ura4-DS/E, (Chl6 ade6-216)* |
| **FY2002** | $h^+$ *leu1-32 ura4DS/E ade6DN/N imr1L::ura4 otr1R::ade6* |
| **PG1672** | *mat1PΔ17::LEU2 mat3-M(RV) ::ade6ori 1 leu1-32 ura-D18 ade6-210* |
| **SPJ1009** | *mat1Msmto leu 1-32 REIIIΔmat3:: ade6 ade6DN/N* |
| **DSPR** | *mat1Msmto leu1-32 ura4D18 REII Δmat2P::ura4 ade6-216 sng2-1(dark derivative)* |
| **LSPR** | *mat1Msmto leu1-32 ura4D18 REII Δmat2P::ura4 ade6-216 sng2-1 (lightderivative)* |
| **SPJ25** | *mat1Msmto leu1-32 ura4D18 ade6-210 his2* |
| **SPP192** | *mat1Msmto leu1-32 ura4D18 REII Δmat2::ura4 ade6-216 with YFP-FLAG-(his)6 tagged cut4 integrated at leu1 by digesting it with ApaI* |
| **SPP91** | *mat1PΔ17::LEU2 REII Δmat2P::ura4 leu1-32 ura4D18* $his2^-$ *ade6-210* $[SNG2]_P$ |
| **SPP79** | *mat1Msmto REIIΔmat2 ura4 leu1-32 ura4D18 ade6-216 clr3Δ::*$kan^r$ *(dark)* |
| **SPP75** | *mat1Msmto leu1-32 ura4D18 REII ΔmatP2::ura4 ade6-216 hsp104Δ::* $kar^r$ |
| **SPP76** | *mat1Msmto leu1-32 ura4D18 REII Δmat2P::ura4 ade6-216 cut4::cut4(GFP::*$kan^r$*)* |
| **SPP87** | *mat1Msmto leu1-32 ura4D18 REII Δmat2P::ura4 ade6-2160* $[SNG2]_{mat1M}$ |
| **SPP180** | *mat1Msmto leu1-32 ura4D18 REII Δmat2P::ura4 ade6-216 tht1Δ::* $kan^r$<br>*mat1Msmto leu1-32 ura4D18 REII Δmat2P::ura4 ade6-210 tht1Δ::* $kan^r$ |
| **SPP153** | *mat1PΔ17::LEU2 REII Δmat2P::ura4 leu1-32 ura4D18* $his2^-$ *ade6-216 tht1Δ::* $kan^r$ |
| **SPP162** | *mat1PΔ17::LEU2 REII Δmat2P::ura4 leu1-32 ura4D18* $his2^-$ *ade6-216 tht1Δ::* $kan^r[SNG2]_P$ |
| **SSP01** | $[SNG2]^O$ derivative generated by overexpression of *cut4* gene in strain *SPP75* |
| **SPP181** | *mat1Msmto leu1-32 ura4D18 REII Δmat2P::ura4 ade6-216 tht1Δ::* $kan^r$ $[SNG2]_M$<br>*mat1Msmto leu1-32 ura4D18 REII Δmat2P::ura4 ade6-210 tht1Δ::* $kan^r cut4^+$ |
| **SPP183** | *mat1PΔ17leu1-32 ura4D18 REII Δmat2P::ura4 ade6-210 tht1Δ::* $kan^r$ $[SNG2]_P$<br>*mat1PΔ17leu1-32 ura4D18 REII Δmat2P::ura4 ade6-216 tht1Δ::* $kan^r$ $cut4^+$ |
| **SPP201** | *mat1Msmto leu1-32 ura4D18 REII Δmat2P::ura4 ade6-210 tht1Δ::* $kan^r$ *swi6Δ::his1*<br>*mat1Msmto leu1-32 ura4D18 REII Δmat2P::ura4 ade6-210 tht1Δ::* $kan^r$ |
| **SPP152** | *mat1Msmto leu1-32 ura4D18 REII Δmat2P::ura4 ade6-210 tht1Δ::* $kan^r$ |

**Table S2. Plasmids used in the study**

| **Plasmid Name** | **Description** |
|---|---|
| pREP3X | *nmt1* promoter with *LEU2* marker S.pombe expression vector |
| pYY463 | *cut4*$^{+}$ gene in pREP41 vector under *nmt1* promoter with *LEU2* gene as selectable marker (high copy expression vector) |
| pYY439 | *cut4*$^{+}$ gene (14kb cosmid clone 813) in pYC11 integration vector with *LEU2* gene as selectable marker |
| pYY463-dB/B | Essentially pYY463 vector with a deletion of fragment 848-3124 between two BglII sites of *cut4* gene |
| pREP41HAN | *nmt1* promoter with *LEU2* marker in expression vector for HA tagging ofproteins in *S. pombe* |
| pREP41HAN Cut4 | *nmt1* promoter with *LEU2* marker in expression vector containing HA-Cut4 |
| pREP41HAN Cut4-dB/B | nmt1 promoter with *LEU2* marker in expression vector having truncated HA-Cut4 with a internal deletion at BglII sites |
| YFH-Cut4 | Full length Cut4 plasmid with YFP-(his)6-FLAG tag at N-terminal of cut4procured form Riken DNA Bioresource Center |

#### SUPPLEMENTARY FIGURE LEGENDS

**Figure S1. Non-Mendelian segregation and mitotic stability of the spo+-ura+ phenotype of the [SNG2] prion form. (A)** Table showing the score of $ura^+$ and $spo^+$ phenotypes in the cross shown in Figure 2. Top panel shows the segregation of $ura^+$:$ura^w$ phenotype among the tetrads generated from the cross. Lower panel shows the score for the different combination of the $ura^+$ and $spo^+$ phenotypes among the $ts^+$ and $ts^-$ segregants of the cross. **(B)** Mitotic stability of the phenotypic states displayed by the segregants of the cross shown in Figure 3B. The states D and L represent the $spo^+$ and $spo^+$ phenotypes and $ura^+$, $ura^-$ and $ura^w$ represent strong growth, no growth or weak growth on medium lacking uracil.

**Figure S2. Mendelian segregation of phenotypes generated by a canonical heterochromatin mutant. (A)** $spo^-$-$ura^-$ phenotype of the segregants generated from a cross between the WT parent strains: Tetrads derived from a cross between WT strains with genotype I (*mat1Msmto leu1-32 REIIΔmat2::ura4 ura4D18)* and genotype II *(mat1PΔ17:: LEU2 leu1-32 REIIΔmat2::ura4 his2 ura4D18),* were replica plated onto indicated plates. The genotypes are mentioned above and below the tetrad dissection panel. **(B)** Co-segregation of $spo^+$-$ura^+$ phenotypes with *clr3Δ* mutation during meiosis: Tetrad analysis of a cross between *clr3Δ::kan$^r$* mutant with genotype I that gives spo+/ura+ phenotype and a wild type strain with genotype II. The genotypes are mentioned above and below the tetrad dissection panel.

**Figure S3. de novo generation of [SNG2] prion-form by high level expression of *cut4* and its non-Mendelian segregation. (A)** Serial dilution spotting assay of WT strain with genotype I, transformed with empty vector, integrating vector containing intact *cut4* gene and truncated copy of the *cut4* gene (DB/B). (B) Results of segregation of spo+/ura+ phenotype in a backcross of the $[SNG2]^o$ derivative obtained after loss of the *cut4* gene on a high copy plasmid, shown in Figure 4D from a strain with genotype I with a wt strain with genotype II.

**Figure S4. Lack of intrachromosomal recombination and interchromosomal segregation in a strain with a homozygous karyogamy mutation *tht1Δ/tht1Δ*.** Strain having linked $leu^+$(*mat1P*-linked)/*his2*$^-$ on chrII along with an *ade6-210* allele on chromosome III was crossed with another *tht1Δ* strain having *mat1*-linked *leu1*$^-$/*his2*$^+$ alleles on chromosome II and *ade6-216* allele on chromosome III and subjected to random spore analysis followed by scoring of the *leu1*, *his2* and *ade6* markers. Results show no cross over between *leu* and *his2* loci and show co-segregation of chr II and III.

**Figure S5. Dominant negative effect of [SNG2] prion-form on silencing at *ade6* reporter inserted at the centromere and *mat3* loci**. **(A)** Schematic representation of the centromere I, showing central element *cnt1*, the inner repeats *imr1L* and *imr1R* and outer repeats *otr1L* and *otr1R*. Also shown are insertion of *ura4* and *ade6* reporter at *imr1R* and *otr1R* repeat region, respectively. WT strains wherein both the reporters are silent, display low growth and red colored colonies on plates lacking uracil or limiting amount of adenine, respectively. **(B)** Organization of the mating type locus, showing *mat1P* locus having a deletion of cis-acting region along with a *LEU2* reporter insertion (*mat1PΔ17::LEU2*) and *mat3*-linked *ade6* reporter. WT strains form red colored colonies on media containing limiting adenine. Loss of silencing in (A) and (B) leads to pink/white phenotype of colonies on adenine limiting plates. **(C)** 5 random putative $prion^+$ segregants having *mat1M* allele labelled "$[SNG2]_M$"1-5 were obtained from the cross of $[SNG2]_P$ with *mat1Msmto* strain (SPJ25) and a normal *mat1Msmto* strain (SPJ25). These were further crossed with a strain containing stable *mat1P* locus and having *ade6* inserted at the *otr1* region of centromere **(A, panel a)** or *mat3* locus **(B, panel b).** Random spores from the crosses (1-5), along with a control *Msmto* strain (SPJ25) were streaked on adenine limiting plate (YE), grown at 30°C for 4 days and photographed.

**Figure S6. Overexpression of *cut4* abrogates silencing at the *ade6* reporter inserted at the *otr1R* locus.** Strain having ade6 insertion at the *otr1R* site on *chrI* was transformed with high copy vector *pREP3* and the same vector containing the *cut4* gene or a copy of *cut4* gene having internal deletion of the *Bgl*II-*Bgl*II region (cut4DB/B). Transformants were streaked on selective plates having low amount of adenine. After 3-4 days' growth at 30$^{o}$C, the colonies were counted and photographed.

**Figure S7. PLAAC analysis of Cut4 sequence.**

Upper panel shows PLAAC score, wherein the HMM is in background state throughout (Black) and never enters PrLD state. However, prediction based on PAPA algorithm shows green underlined regions, while regions predicted by Foldindex as intrinsically disordered regions (IDRs) are underlined in black (lower panel).

**A**

| Ratio ura$^+$ : ura$^w$ | No. tetrads |
|---|---|
| 4:0 | 9/ 54 (16%) |
| 3:1 | 12/ 54 (22%) |
| 2:2 | 23/ 54 (41%) |
| 1:3 | 6/ 54 (11%) |
| 0:4 | 4/ 54 (7%) |

| | |
|---|---|
| Total ascospores | 216 |
| ura$^+$ | 124:<br>63 his+<br>61his- |
| ura$^-$ | 92 |
| ts+ ura+ | 64 (30%) |
| ts+ ura- | 40 (18%) |
| ts- ura+ | 60 (29%) |
| ts- ura- | 52 (23%) |
| ts+ spo+ | 34 (16%) |
| ts+ spo- | 70 (32%) |
| ts- spo+ | 47 (22%) |
| ts- spo- | 65 (30%) |

B

| | Nature of switch | rate of switching of spo phenotype |
|---|---|---|
| DSPR | D to L | $2.7X10^{-4}$ |
| LSPR | L to D | $1.9X10^{-4}$ |
| Msmto REIIΔmat2::ura4 | L to D | $1.05X10^{-5}$ |
| [SNG2] spo$^+$-ura$^+$ | D to L | $5.6X10^{-5}$ |
| [SNG2] spo$^-$-ura$^+$ | L to D | $1.7X10^{-4}$ |
| [SNG2] spo$^+$-ura$^w$ | D to L | $2.3X10^{-5}$ |
| [SNG2]$_M$ spo$^w$-ura$^+$ | L to D | $2.1x10^{-4}$ |

| | Nature of switch | rate of switching of ura phenotype |
|---|---|---|
| DSPR | ura$^+$ to ura$^-$ | $2.8X10^{-4}$ |
| LSPR | ura$^+$ to ura$^-$ | $3.9X10^{-4}$ |
| Msmto REIIΔmat2::ura4 | ura$^-$ to ura$^+$ | $2.8X10^{-3}$ |
| [SNG2]ura$^+$spo$^+$ | ura$^+$ to ura$^-$ | $3.2X10^{-4}$ |
| [SNG2]ura$^+$spo$^-$ | ura$^+$ to ura$^-$ | $1.00X10^{-4}$ |
| [SNG2]ura$^w$spo$^+$ | ura$^w$ to ura$^+$ | $9X10^{-5}$ |
| [SNG2]M ura$^+$spo$^w$ | ura$^+$ to ura$^-$ | $9.5X10^{-6}$ |

Supplementary Figure S1

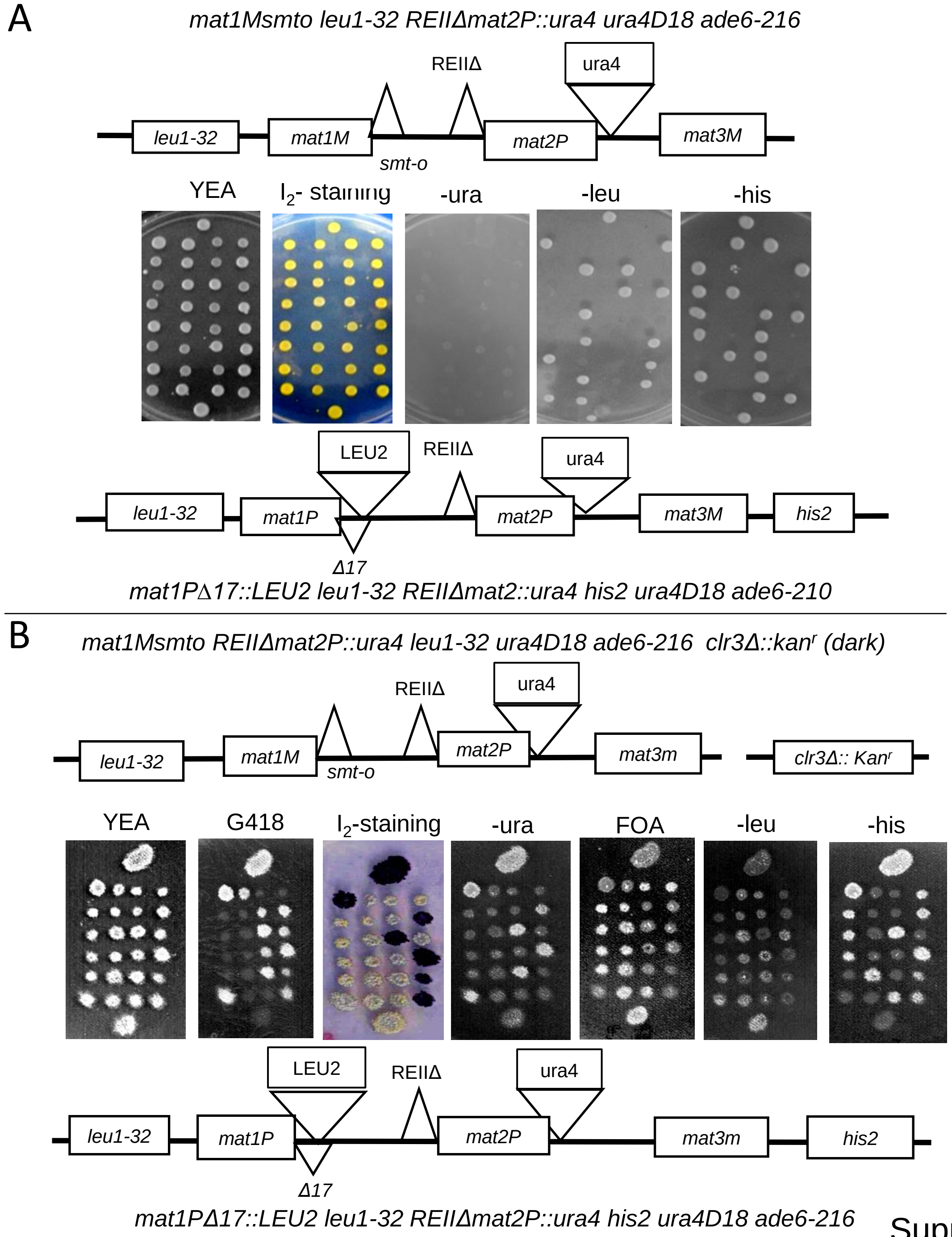


Supplementary Figure S2

**A**

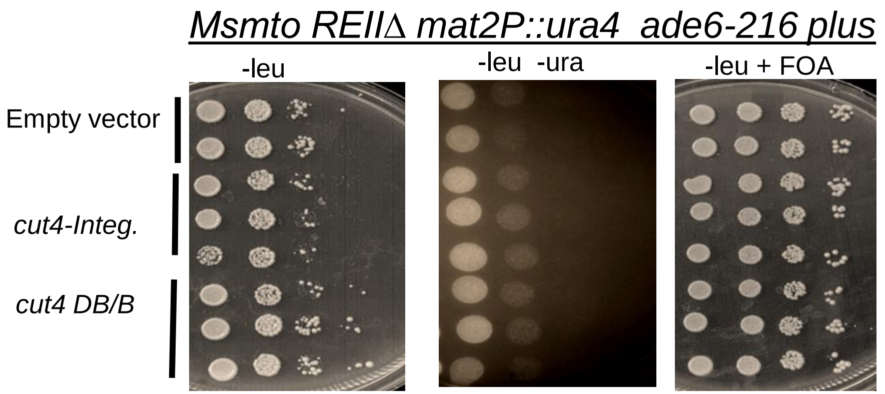


**B**

| Ratio of $ura^+$: $ura^w$ | No. of tetrads (%) | $ura^+$ $leu^+$ $his^+$ | $ura^+$ $leu^-$ $his^+$ | $ura^+$ $leu^+$ $his^-$ |
|---|---|---|---|---|
| 4:0 | 7/50 (14) | 8 | 12 | 8 |
| 3:1 | 29/50(58) | 26 | 40 | 21 |
| 1:3 | 1/50 (2) | 1 | 0 | 0 |
| 2:2 | 13/50 (26) | 6 | 10 | 10 |

Supplementary Figure S3

| *Msmto leu1-32 ura4D18 REII Δmat2P::ura4 ade6-216 tht1Δ::kan$^{r}$*<br>X<br>*PΔ17::LEU2 REII Δmat2P::ura4 leu1-32 ura4D18 his2$^{-}$ ade6-210 tht1Δ::Kan$^{r}$*<br>(**Total no. of spores N= 276**) | | | |
|---|---|---|---|
| leu$^{+}$ -his$^{-}$ | 117 | *ade6-210* | 117 |
| | | *ade6-216* | 0 |
| leu$^{--}$-his$^{+}$ | 159 | *ade6-210* | 0 |
| | | *ade6-216* | 159 |
| leu$^{-}$ -his$^{-}$ | 0 | - | |
| leu$^{+}$-his$^{+}$ | 0 | - | |

Supplementary Figure S4

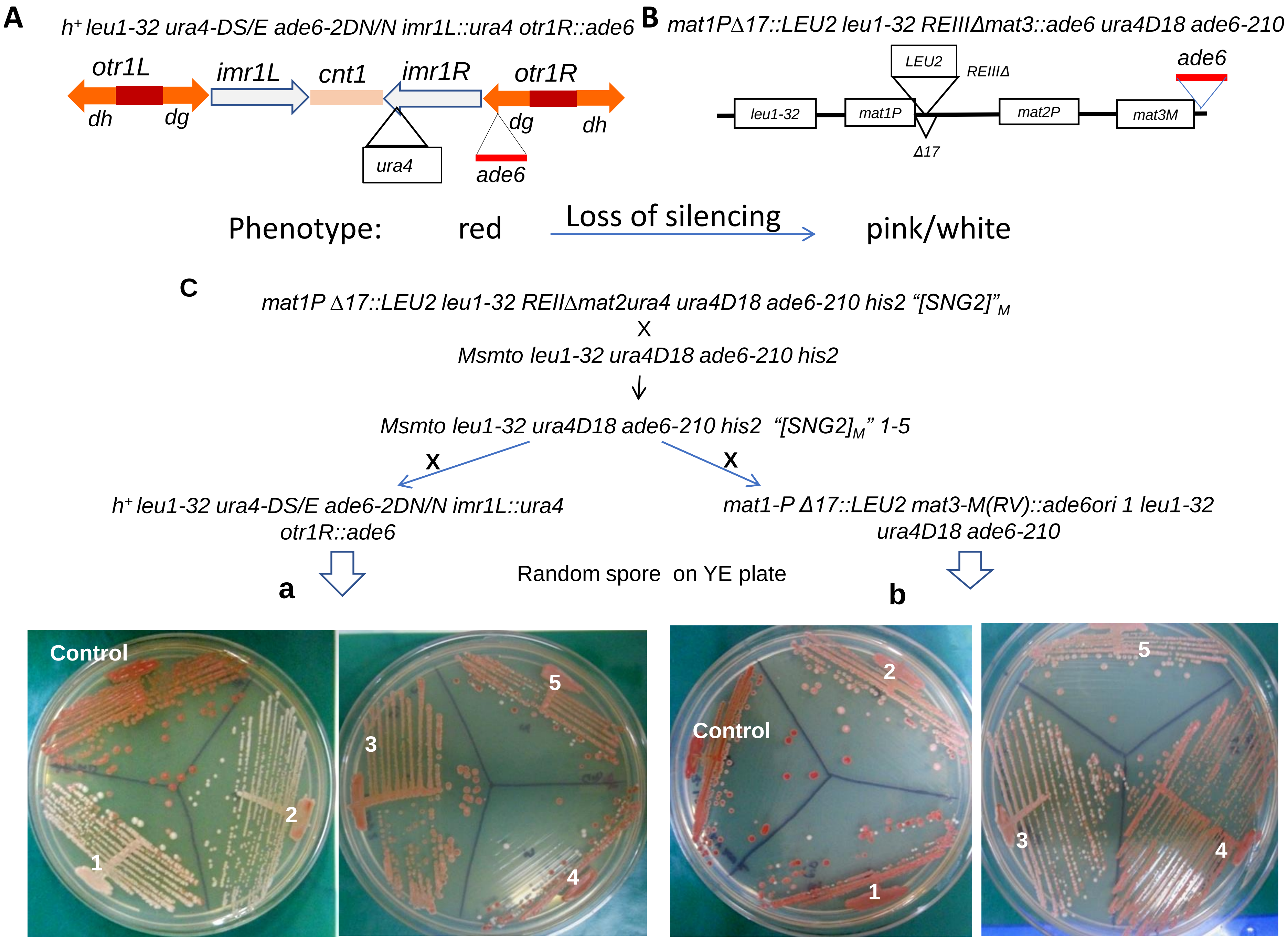


Supplementary Figure S5

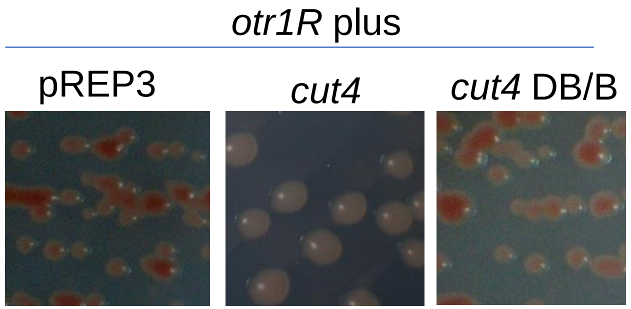


| | pREP3 | *cut4* | *cut4* DB/B |
|---|---|---|---|
| % Pink | 11 | 35.4 | 19.8 |
| % white | 1.9 | 39.2 | 0 |

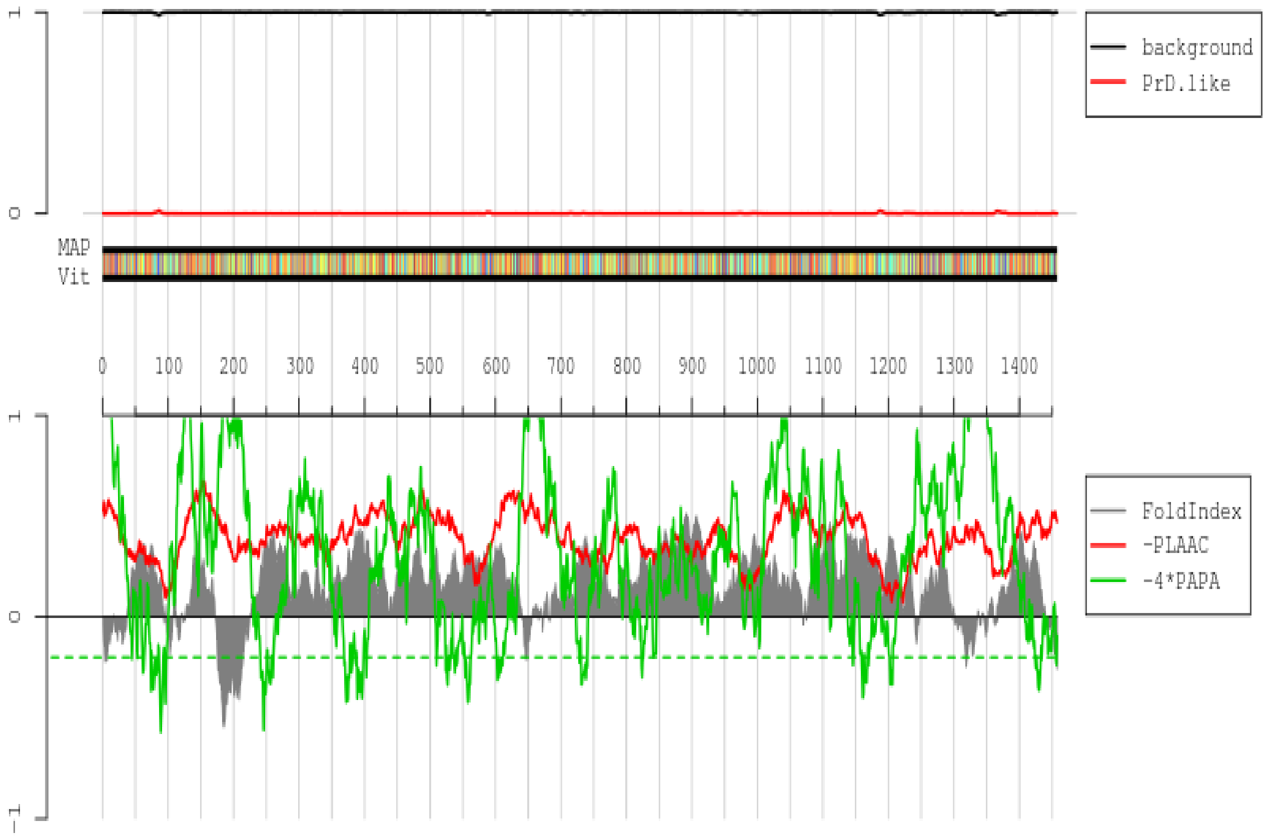


1 MLELKTNVIQDPELKDSLDLQENDRVEIIGDAVHYIVNDHLNRVFNYTVDQQKIHAALITTFASGKKAIVVILDDIGYVYYVGDNNNDSYIINVPFSTES
101 AWSSPSGLYLQRHRSSDENLNTDLPHIFCLNDPLDELTLIKFDGKKILSLFDSIVYVVGEIVVTHNKKEKKLSFWRSRFIDPESDQSIKSSRNRRRESSF
201 SREKNPDLTRSDSIHYTANTRLSEKFEEQGLAYSTIFSHIESFPITGSTSFDSILGNGVLVITTLVKELEKGYMMLFRLVRDRSPYFLDSLQLHAINLSA
301 IKSKHLQKIVVLSSKGKVSLESPMSPSLPIEGTFRSFRVHGATLYLEDTDGVQRYISLDNRASNSLVKWCLSVIRYVLPLREYEIFYTGHLYALFAFKLS
401 HDEAFISSILACFTFFSRDKVHVEPIEDCNEAYSLSSKFHFKKEILIASQLSSHLDYSTFKNYLMPLAITLHFISEELRLDSVVKPRKDQLVALLLQITT
501 WLKWPRYCEYYNFDIAETFLSIPLSIQVDVEEPVGPTSILQWIIECLRSQSTVPFYGLESYGLPHSCSTMFPQTLSLMQLLDCLLNPNMTLQNLVEEMVR
601 LGISRKRCERYPFGILCIIFTVLEIAAEEYSPNWESEELRLVNRLDVDSFLHPKTPKWVFNKQDQEVKEIKALTSTVTDSTLVDTQSFHPYKVVTDMIFR
701 EDRRLAEVNKLLNYSSQITIMTEHFDVDLSSVPMQQKVAQCICVRTLSVPIGAGMLTYGSKNPLPTEKVTPRLFNFTLHLHPGTLIIQPNKEFVTQELTE
801 WPEFNVGVALGLSISKFSKEINTSWIMFNRPETLTAYHAGFLFGLGLNGHLKALATWHSFIYLTSKHDTTSIGLLLGLASSYLGSMDAKVTKLLSVHISA
901 LLPVGSNELNISPLTQTAGILGIGLLFHDSCHRRMSEVTMEEILASNESELKNEGYKLAAGFSLGLINLGRGSNLPGMSDLKLVSRLQVGISSQATFQSL
1001 EAGSPGAIMALTMIYMKTNDLEVAKKIDIPKSRYLLDFYRPDLILLRVAGKNLIMWDEVKADYEWVKYQIPDIMLSQFDLQEKKVLSSDDLLLYNVLAGI
1101 CFSLGLRFAGTGNPKAKEILINFLDSFIRLCHLPAKTHDERVTAVTVIRCTQIVALSSSCVMAGYCDLDVLRRLRVLHGRMEPVNYGAQMATHMALGILS
1201 LGGGRYSLSRSNLAIAALLISFYPQFPRTTQDNRAHLQAARNLWALAVEERCIIPRNQDTKQPCIVPLNVVQKSGAVQKLEAPILLPPYDSISSVSTLGD
1301 KYWNLKIDLDNNSDYRELLRESQTLTLMPYDRTSSKEEPLNLFPKLKDTSSPLWNLVKTSRLFQSSNSPLNVASLQESNNKTSLGVKLLLSMDFDNLTRD
1401 RLLSLQILLQFFESCWTGVLLNKFHSRQYLFLSRDLVEDLSLRVWEYVYSHNHNEESV

Supplementary Figure S7
